# A direct interaction between the RNA-binding proteins Staufen and Tm1-I/C regulates *oskar* mRNP composition and transport

**DOI:** 10.1101/2024.12.18.629124

**Authors:** Thomas Gaber, Julia Grabowski, Bernd Simon, Thomas Monecke, Tobias Williams, Vera Roman, Jeffrey Chao, Janosch Hennig, Anne Ephrussi, Dierk Niessing, Simone Heber

## Abstract

In the *Drosophila* female germline, *oskar* messenger RNA is transported on microtubules from the nurse cells to the posterior pole of the oocyte, where it is translated. Transport of *oskar* transcripts from the nurse cells into the oocyte requires dynein, while localization of the mRNAs within the oocyte to the posterior pole is dependent upon kinesin-1. Staufen, a dsRNA-binding protein, has been shown to bind the *oskar* mRNA transport complex in the oocyte and inactivate dynein; however, it remains unclear how kinesin is activated. Here, using surface plasmon resonance, nuclear magnetic resonance spectroscopy and RNA imaging within egg chambers, we demonstrate that Staufen directly interacts with Tm1, a non-canonical kinesin adaptor. This work provides a molecular explanation for the previously unclear role of Staufen in *oskar* mRNA localization.

## Introduction

Active transport of organelles, vesicles and mRNAs by mechanoenzymes along microtubules is essential for the functional polarization of cells. Such cellular cargoes are often associated with both dynein and kinesin – motor proteins of opposite polarity – and, as a consequence, travel bidirectionally (Welte, 2004; Hancock, 2014). Coordination of the motor protein activities is therefore required to avoid a futile tug-of-war. Both dynein and kinesin adopt autoinhibited states, and activation is driven by interactions with adaptor proteins, cargoes and microtubule binding proteins (Reck-Peterson *et al*, 2018; Olenick & Holzbaur, 2019; Verhey & Hammond, 2009; Chiba *et al*, 2022).

o*skar* mRNA localization in the *Drosophila* egg chamber is a paradigm for the study of RNA transport and motor protein regulation during dual-motor transport. *oskar* localization during oogenesis drives abdominal patterning and germline formation in the embryo, and mislocalization of the mRNA has profound phenotypic consequences (Ephrussi & Lehmann, 1992). In the germline syncytium, *oskar* mRNA synthesized in the nurse cells is assembled into messenger ribonucleoprotein (mRNP) complexes and actively transported into the transcriptionally silent oocyte (Laver *et al*, 2015; Roth & Lynch, 2009; Wilhelm & Smibert, 2005) by the minus-end directed motor dynein in complex with dynactin and the activating adaptor Bicaudal D (BicD) (Clark *et al*, 2007). The RNA-binding protein Egalitarian (Egl) links the *oskar* transcript to the transport complex via a stem-loop structure, called oocyte entry signal (OES), in the *oskar* 3’UTR (Jambor *et al*, 2014; Clark *et al*, 2007; Bullock & Ish-Horowicz, 2001; Dienstbier *et al*, 2009; Navarro *et al*, 2004). In the oocyte, the plus-end directed motor kinesin-1 (kinesin heavy chain, Khc) takes over and transports *oskar* along the polarized microtubule cytoskeleton to the posterior pole (Brendza *et al*, 2000; Zimyanin *et al*, 2008), where the mRNA is translated (Rongo *et al*, 1995). This Khc-mediated transport relies on the cis-acting spliced *oskar* localization element (SOLE), which forms upon splicing of the first *oskar* intron, and deposition of the exon junction complex (Hachet & Ephrussi, 2004; Ghosh *et al*, 2012) on the mRNA.

Throughout its transport, *oskar* mRNPs are associated with both the dynein and kinesin transport machineries. Thus, translocation from the nurse cells to the oocyte posterior pole requires motor switching and precise spatial and temporal control of the two motor activities. After translocation of *oskar* mRNPs into the oocyte, dynein is antagonized by recruitment of the double-stranded RNA-binding protein (dsRBP) Staufen. This causes dissociation of the adaptor protein Egl from the mRNA (Gáspár *et al*, 2023; Mohr *et al*, 2021) and the mRNP switches to kinesin-1 (kinesin heavy chain, Khc)-mediated transport. The atypical I/C-isoform of Tropomyosin-1 (Tm1) functions as an RNA cargo adaptor (Gáspár *et al*, 2017; Veeranan-Karmegam *et al*, 2016; Dimitrova-Paternoga *et al*, 2021), both linking Khc to the mRNA and inhibiting Khc activity during dynein-mediated transport. Tm1 promotes autoinhibitory interactions within the Khc stalk domain via a mechanism that involves its N-terminal domain (Heber *et al*, 2024). How Khc is activated in the oocyte remains unclear. Because Tm1 is associated with the mRNP during Khc-mediated transport, Khc activation likely involves remodeling of the mRNP and a conformational change of the Khc-Tm1 complex. Staufen, which associates with the *oskar* mRNP upon entry into the oocyte and displaces Egl, is a central factor of mRNP remodeling. It is also a likely inducer of motor switching.

To elucidate the *oskar* mRNP remodeling events underlying motor switching, we screened known proteins involved in Khc-mediated *oskar* localization (tubulin, exon junction complex, Khc, Tm1) for a direct interaction with Staufen. Using surface plasmon resonance (SPR) and nuclear magnetic resonance (NMR) spectroscopy, we show that Staufen interacts directly with Tm1. We map the binding sites in both interaction partners and show by mutational analysis that the interaction between Staufen and Tm1 can be specifically abolished. By imaging of *oskar* RNA and transgenic wild-type and mutant Tm1 proteins in *Drosophila* egg-chambers, we show that loss of the Staufen-Tm1 interaction causes mislocalization of Tm1 *in vivo* and partially impairs *oskar* localization, demonstrating the importance of the Staufen-Tm1 interaction for *oskar* mRNP transport. These findings indicate that, in addition to inhibiting dynein, Staufen also plays a role in Khc activation, coupling regulation of the two microtubule motors.

## Results

### Staufen and Tm1 interact directly

To identify direct interactions between Staufen (Figure 1A) and components of the *oskar* mRNA localization machinery, we performed surface plasmon resonance (SPR) experiments with recombinant proteins (Figure S1). Several proteins known to be involved in *oskar* mRNA transport within the oocyte were injected as analytes to test for direct interactions with immobilized Staufen in single-cycle experiments.

**Figure 1:**
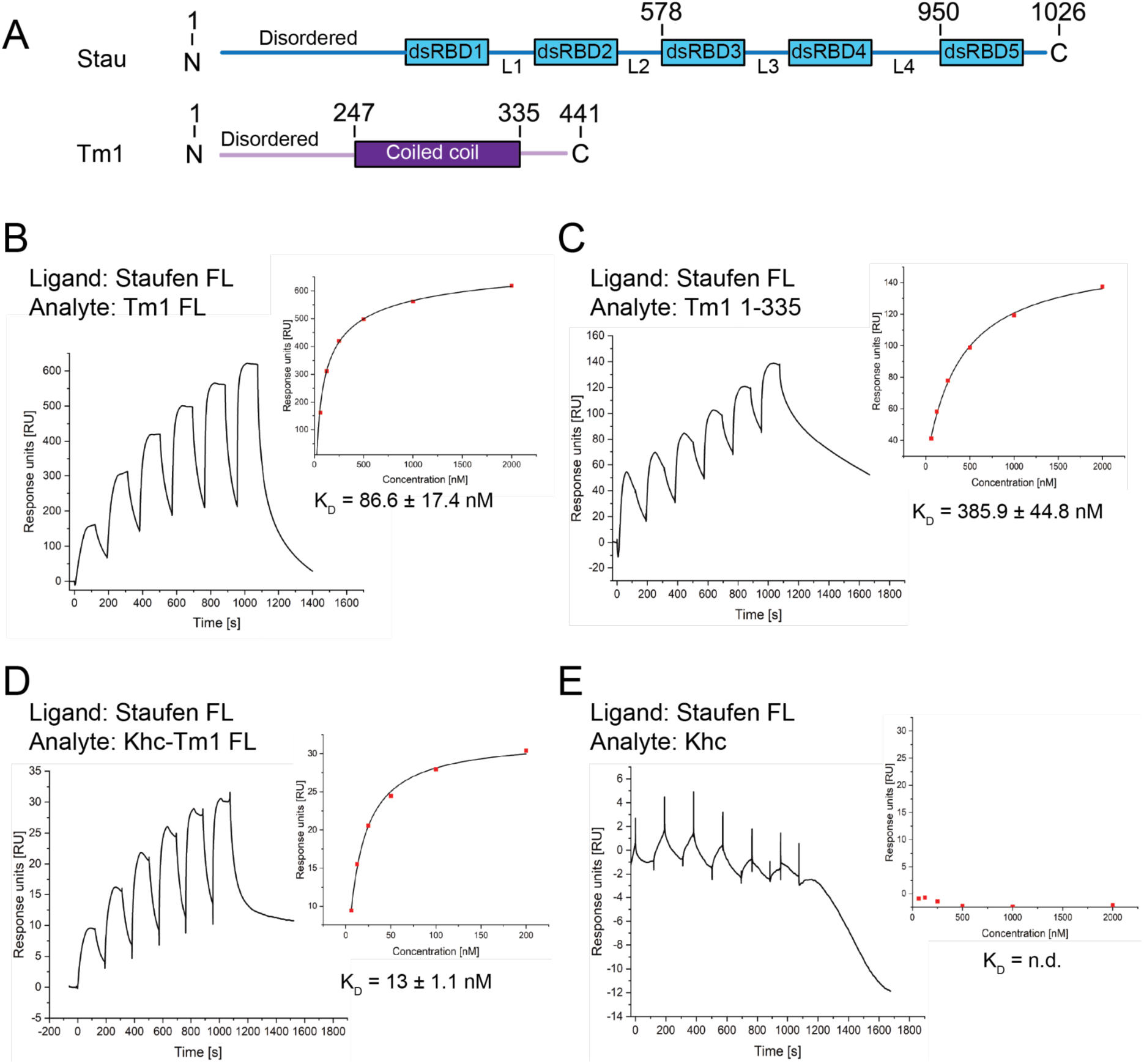
Recombinant Staufen and Tm1 interact directly and form a ternary complex with Khc. **A**) Scheme of *Drosophila melanogaster* Staufen and Tm1 proteins. dsRBD = double-stranded RNA-binding domain, L = linker. **B-E)** SPR experiments show direct binding of Staufen FL to Tm1 FL (B), to the truncated N-terminal construct Tm1 1-335 (C), and to the Khc-Tm1 complex (D). No binding of Staufen FL to Khc was observed (E). For these experiments, GST-Staufen FL was surface-coupled by GST-capture with an anti-GST antibody. The sensorgrams of single-cycle SPR experiments (left in each panel) show that Tm1 proteins bind transiently to Staufen. Equilibrium dissociation constants (K_D_) were determined using a one-site-binding fit to the maximal response at equilibrium for each analyte concentration. K_D_-values are given as mean ± standard deviation (SD) from triplicate experiments. Concentrations of analytes used in (B), (C), and (E) were 62.5, 125, 250, 500, 1,000, and 2,000 nM. Concentrations of analyte used in (D) were 6.25, 12.5, 25, 50, 100, 200 nM.

While we could not detect an interaction with the core components of the exon junction complex (Btz-1-345-eIF4AIII and Mago-Y14 heterodimers (Bono *et al*, 2006; Le Hir & Andersen, 2008; Andersen *et al*, 2006), Figure S2A,B) or the cytoskeletal protein tubulin (Figure S2C), we observed an interaction of the kinesin cargo adaptor Tm1 with the surface-coupled Staufen protein (Figure 1A). The observed binding event was transient with an equilibrium dissociation constant (K_D_) of 86.6 nM –± 17.4 nM (Figure 1B).

We previously showed that an N-terminal fragment of Tm1 (aa 1-334) is sufficient to rescue Tm1’s function during *oskar* mRNA localization in *Drosophila* oocytes (Dimitrova-Paternoga *et al*, 2021). Thus, we tested whether Tm1 1-335 is also sufficient for the interaction with Staufen. In comparison to the full-length protein (Tm1 FL), Tm1 1-335 exhibited a slightly weaker interaction, with a K_D_ of 385.9 nM ± 44.8 nM and similar on and off rates (Figure 1C).

In the *Drosophila* ovary, Tm1 forms a complex with Khc. We therefore tested if Khc-bound Tm1 interacts directly with Staufen in a ternary complex. Indeed, co-purified Khc-Tm1 bound to Staufen with a K_D_ of 13 nM ± 1.1 nM (Figure 1D), whereas Khc alone did not exhibit any interaction with Staufen (Figure 1E). This result shows that the interaction between Staufen and Tm1 is not impeded by Tm1’s interaction with Khc, suggesting that Tm1 might serve as an adapter between Staufen and Khc.

### Mapping the interaction between Staufen and Tm1

To better understand how Staufen and Tm1 interact, we sought to identify the specific binding interface between the two proteins.

Staufen consists of 1026 amino acids containing an N-terminal intrinsically disordered region (IDR) and five structured double-stranded RNA-binding domains (dsRBDs) connected by intrinsically disordered, low-complexity linkers (Figure 1A).

To map the Staufen-Tm1 interaction, we purified truncated recombinant Staufen proteins and tested them for direct interaction with surface-coupled Tm1 FL using SPR. The truncated N-terminal Staufen N-term-dsRBD1-2 (Figure S3A, Table S1) did not interact with Tm1 FL, indicating that neither the N-terminal unstructured region nor the dsRBDs 1 and 2 contribute to Tm1 FL binding. A second fragment comprising dsRBDs 3, 4, 5 and the disordered C-terminus (dsRBD3-4-5-C-term) bound to Tm1 FL with a K_D_ of 152 nM ± 60 nM (Figure S2B, Table S1). This equilibrium dissociation constant is similar to the K_D_ measured for binding of Tm1 FL to surface-coupled full-length Staufen (Staufen FL, Figure 1B), showing that the region of Staufen comprising dsRBD3-4-5-C-term is sufficient for binding to Tm1. We also tested Staufen dsRBD1-4, Staufen L4-dsRBD5-C-term, and Staufen dsRBD5-C-term (Figure S3C,D,E, Table S1), but none of these constructs showed binding to Tm1 FL at concentrations up to 2 µM.

Based on these insights, we purified Staufen dsRBD3-4-L4 (aa 578-951) which contains dsRBDs 3 and 4, as well as the intrinsically disordered linker region between dsRBDs 4 and 5. To test if this construct harbors the necessary regions for interaction with Tm1, Staufen dsRBD3-4-L4 was injected onto a CM5 sensorchip with surface-coupled Tm1 1-335. The two proteins interacted transiently with a K_D_ of 224 nM ± 21.46 nM (Figure S4A). The reverse experiment, testing the interaction of Tm1 1-335 to surface-coupled Staufen dsRBD3-4-L4, resulted in a comparable K_D_ of 262.3 nM ± 15.5 nM (Figure S4B).

We then asked whether Tm1 interacts with Staufen via Tm1’s N-terminal IDR or via the coiled-coil domain (Figure 1A). When testing a construct comprising only the N-terminal Tm1 IDR (aa 1-213) we found an interaction with Staufen dsRBD3-4-L4 with a K_D_ similar to Tm1 1-335 (494.8 nM ± 83.9 nM) (Figure 2A). Based on these data, we conclude that the interacting regions of Tm1 and Staufen are located within aa 1-213 of Tm1 and within the dsRBDs 3 and 4 or the linker 4 region of Staufen.

**Figure 2:**
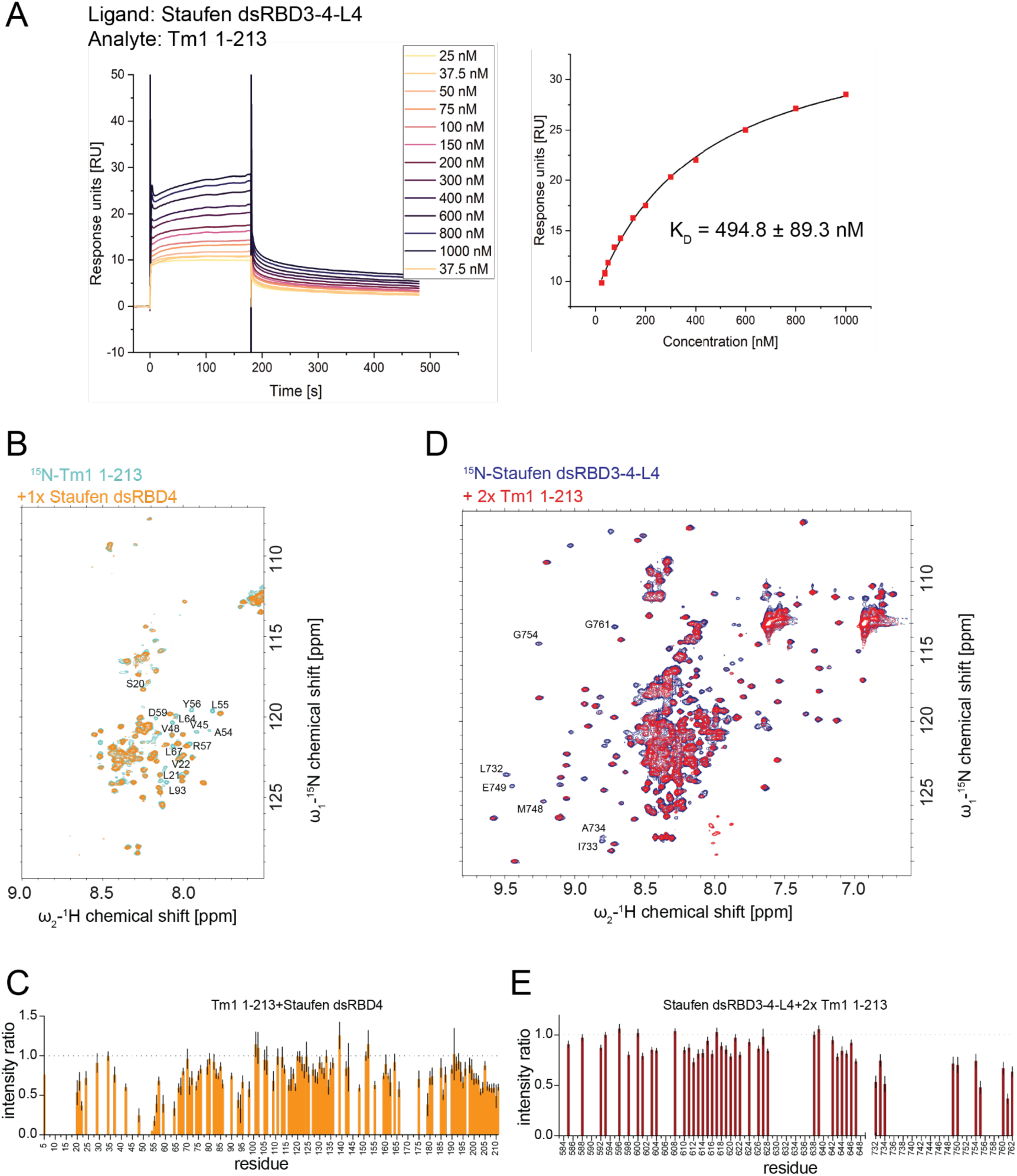
Staufen and Tm1 interact via hydrophobic residues in Staufen dsRBD4 and the Tm1 N-terminal domain. **A**) Tm1 1-213 binding to surface-coupled Staufen dsRBD3-4-L4. The K_D_-value was determined by SPR measurements from the response concentration plot (right) to be 494.8 nM. The corresponding sensorgram is displayed on the left. Twelve individual analyte concentrations were used. To account for possible baseline shifts, one sample concentration was measured in duplicate. The experiment was carried out in triplicate and the K_D_-value determined from one-site binding fits ± SD. **B, C)** NMR titrations of ^15^N-Tm1 1-213 with Staufen dsRBD4. Overlay of ^1^H-^15^N HSQC spectra of free Tm1 1-213 (cyan) and Staufen dsRBD4-bound Tm1 1-213 (orange). Relevant backbone assignments (Vaishali *et al*, 2021) are indicated (B). Plotted intensity ratios of individual residues in the titration experiment (C). **D, E)** NMR titrations of ^15^N-Staufen dsRBD3-4-L4 with Tm1 1-213. ^1^H-^15^N HSQC spectra of free Staufen dsRBD3-4-L4 (blue) with Tm1 1-213-bound Staufen dsRBD3-4-L4 (red). Relevant backbone assignments are indicated (D). Plotted intensity ratios of individual residues in the titration experiment (E). Data are presented as measured value ± SD. Error bars represent estimates of propagated measurement errors of the experimental uncertainties in signal amplitudes.

### Determining the Tm1 and Staufen interaction at amino-acid resolution

Having identified the domains involved in the interaction between Tm1 and Staufen, we now aimed to determine their interaction interface at amino acid resolution. To this end, we performed nuclear magnetic resonance (NMR) titration experiments.

First, we acquired ^1^H-^15^N HSQC spectra for Tm1 1-213 (Vaishali *et al*, 2021) with increasing amounts of unlabeled Staufen dsRBD3-4-L4. Staufen decreased the intensity of resonances corresponding to several residues in the N-terminal region of Tm1 (Figure S6A,B), indicating a direct interaction due to the decreased molecular tumbling of a larger complex. Small chemical shift perturbations of the resonances of several amino acids (Figure S6C) further confirm the interaction.

Staufen dsRBD3-4-L4-binding affected resonance intensities for a large region of the Tm1 N-terminus, likely due to line broadening upon binding of a relatively large interaction partner. Therefore we repeated the titration experiment with the individual Staufen dsRBDs 3 and 4 to identify residues directly involved in the interaction, whose peak intensities are most strongly affected. While fragments smaller than dsRBD3-4-L4 did not interact detectably with Tm1 in the SPR experiment, NMR experiments are performed at much higher concentrations (one to two orders of magnitude) and might detect even weaker interactions. We thus reasoned that we could identify the amino acids most affected by the interaction. Using this approach, we found that both dsRBD3 and dsRBD4 affected N-terminal residues in Tm1 1-213 (Figure 2B, C; Figure S7A,B), yet dsRBD4 showed a much stronger effect at a 1:1 ratio between the two proteins than dsRBD3 even at an 8-fold excess (Figure 2E, Figure S7B). This indicates that Tm1’s interaction with dsRBD4 is specific, whereas the interaction with dsRBD3 is due to charge-charge interactions which occurr unspecifically at very high concentrations and are likely irrelevant *in vivo*.

Among the amino acids involved in the binding of Tm1 1-213 to Staufen dsRBD4, the strongest effects were observed for S20, L21, V22, V45, V48, A54, L55, Y56, D59, L64, L67, and L93, indicating a hydrophobic interaction between Staufen and Tm1.

The reverse experiment, titrating Tm1 1-213 to ^15^N-Staufen dsRBD3-4-L4 resulted in strong intensity decreases for several Staufen resonances, indicating that the corresponding amino acids are involved in the interaction or close to the interaction surface. Residues E731, L732, I733, V747, M748 and E749 in the β-sheets of dsRBD4 (Figure S8A) showed the strongest intensity decreases (Figure 2D,E). Four of these amino acids possess a hydrophobic side chain, consistent with a hydrophobic interaction between the two proteins.

### Structure-guided mutations disrupt the interaction between Staufen and Tm1

To validate our findings, we generated point mutations, substituting amino acids which showed considerable changes in the chemical environment upon binding in NMR experiments.

The selected mutations in Tm1 were S20A, L21D, V22T, V45Y, V48S, A54S, L55D, Y56S, D59L, L64E, L67T, and L93Y, mostly replacing hydrophobic by hydrophilic side chains. The mutations chosen for Staufen were E732R, L733T, I734Y, V748F, M749S and E750V. We verified by NMR that these mutations do not alter the folds (or lack thereof in case of Tm1 1-213) of the mutated proteins, comparing their ^1^H-^15^N HSQC spectra to those of the wild-type (WT) proteins (Figure S8B,C, Figure S9).

We first tested the Tm1 mutants for binding to surface-coupled Staufen dsRBD3-4-L4. Whereas wildtype Tm1 1-213 (Tm1 1-213 WT) bound to dsRBD3-4-L4 with a K_D_ of of 494.8 nM ± 83.9 nM (Figure 3A), the mutant Tm1 protein carrying all 12 mutations (Tm1 1-213 M12) showed no saturation of the interaction in the identical experiment (Figure 3B, Table S1). This indicates that the mutated protein does not bind to Staufen dsRBD3-4-L4 at concentrations up to 2 µM.

**Figure 3:**
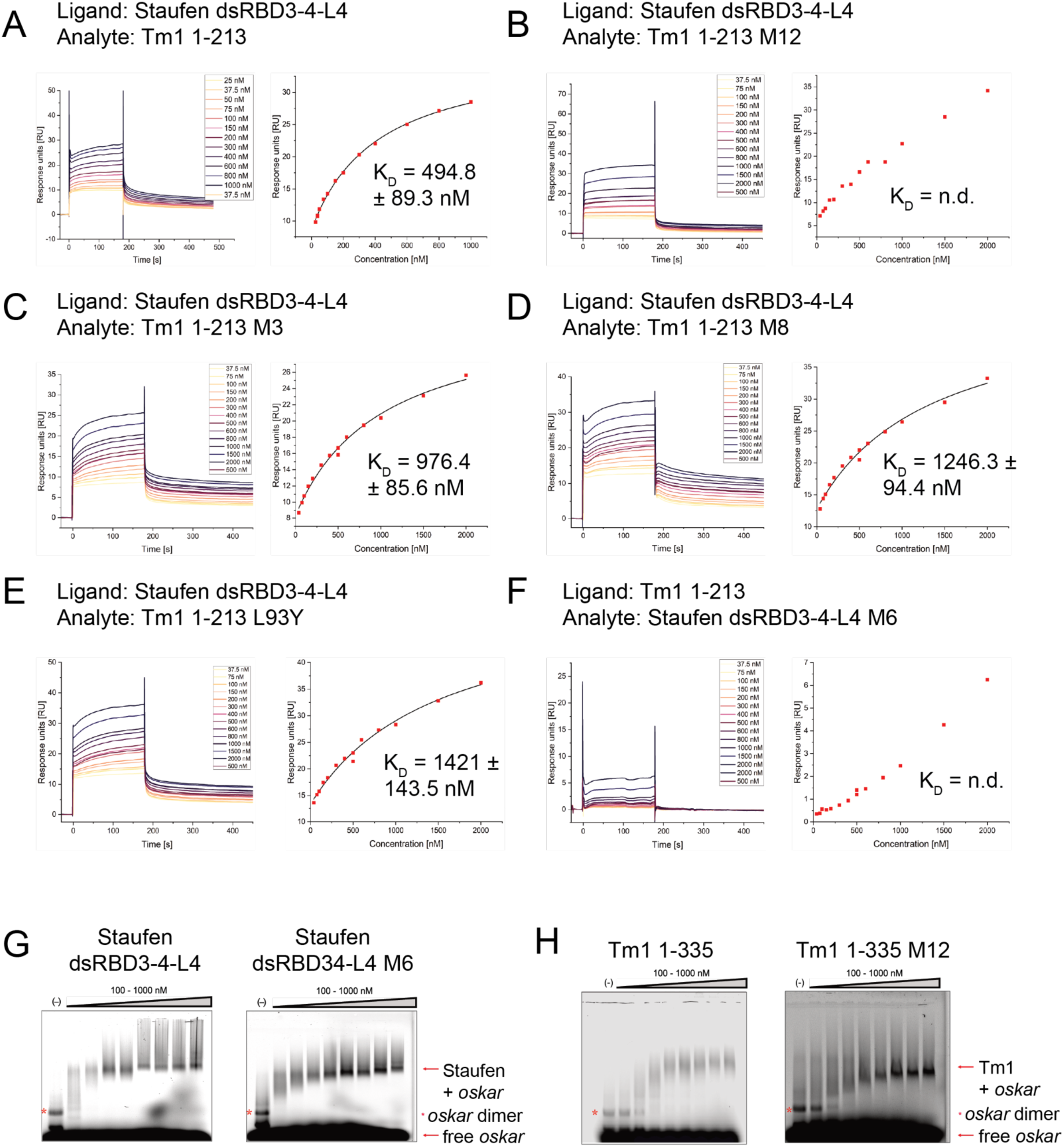
The interaction between Staufen and Tm1 is specifically weakened or abolished by mutations in their binding interface without affecting their RNA-binding properties. Sensorgrams, response concentration plots and determined K_D_s of binding to Staufen dsRBD3-4-L4 by **A)** Tm1 1-213 WT (K_D_: 494.8 nM), **B)** Tm1 1-213 M12 (no saturation), **C)** Tm1 1-213 M3 (K_D_: 976.4 nM), **D)** Tm1 1-213 M8 (K_D_: 1246.3 nM), or **E)** Tm1 1-213 L93Y (K_D_: 1421 nM). **F)** Binding of Staufen dsRBD34-L4 M6 to Tm1 1-213 (no saturation). Wildtype or mutant Tm1 constructs were applied to surface-coupled Staufen dsRBD3-4-L4. Staufen dsRBD3-4-L4 M6 was applied to surface coupled Tm1 1-213. Each experiment was conducted with 13 individual analyte concentrations. To account for possible baseline shifts, one sample concentration was measured in duplicate. K_D_-values were computed by one-site binding fits. Experiments with given K_D_-values were conducted as triplicates. Data are presented as measured value ± SD. Data for Tm1 1-213 WT are reproduced from Figure 2A. **G)** EMSA with dsRBD3-4-L4 WT or M6; **H)** Tm1 1-335 WT or M12. Proteins were incubated with *oskar* 3’UTR region 2+3 (Jambor *et al*, 2014; Gáspár *et al*, 2023) fluorescently labeled with aminoallyl-UTP-ATTO-680. Free RNA and protein-RNA complexes are marked with arrows. All proteins were titrated according to the following concentration series: 100, 200, 400, 500, 600, 700, 800, 1000 nM.

To narrow down the effects of Tm1 mutation on Staufen binding, we divided the mutated region into an N-terminal region carrying three mutations (Tm 1-213 M3), a middle region carrying eight mutations (Tm1 1-213 M8), and a C-terminal region carrying a single mutation (Tm1 1-213 L93Y). When we performed SPR experiments with surface-coupled Staufen dsRBD3-4-L4, we observed two-to three fold reduced binding for all three mutant versions of Tm1 (K_D_s for Tm1 1-213 M3: 976.4 nM ± 85.6 nM, Tm1 1-213 M8: 1246.3 nM ± 94.9 nM, and Tm1 1-213 L93Y: 1421 nM ± 143.5 nM) when compared to wildtype Tm1 1-213 (Figure 3D-E, Table S1). In contrast to the Tm1 M12 mutant, mutations in only one of the three regions did not completely abolish the interaction with Staufen, but weakened it significantly. This observation suggests that all three Tm1 regions contribute to Staufen binding.

When we tested the binding of Staufen dsRBD3-4-L4 harboring all six mutations (Staufen dsRBD3-4-L4 M6) to surface-coupled Tm1 1-213, we failed to observe a saturation in binding with sample concentrations as high as 2 µM (Figure 3F). This observation confirms that the six amino acid residues are required for the interaction between Tm1 and Staufen dsRBD3-4-L4.

To detect any remaining weaker interactions between Staufen and the Tm1 M12 mutant that might occur at higher concentrations than can be assayed by SPR, we performed NMR titrations. When titrating Staufen dsRBD3-4-L4 against ^15^N-Tm1 1-213 M12, we observed no changes in the Tm1 1-213 M12 ^1^H-^15^N-HSQC spectrum, even with an excess of Staufen dsRBD3-4-L4 (Figure S10). This shows that the two proteins do not interact even at high micromolar concentrations (50 µM).

To evaluate whether the *in vitro* tested mutations in Tm1 or Staufen dsRBD3-4-L4 alter the RNA-binding capabilities of the two proteins of interest, we compared the mutated and wildtype proteins in EMSAs with a 534 nt fragment of the *oskar* 3’UTR previously shown to support *oskar* transport by dynein (*oskar* 3’UTR region 2 + 3 (Jambor *et al*, 2014; Gáspár *et al*, 2023)). Staufen dsRBD3-4-L4 and the mutant Staufen dsRBD3-4-L4 M6 (Figure 3G) bound *oskar* at comparable concentrations with apparent K_D_s of ∼400 nM. Tm1 1-335 and the mutant Tm1 1-335 M12 (Figure 3H) also bound RNA at comparable concentrations in the nanomolar range (apparent K_D_ ∼600 nM). This suggests that the selected mutations do not affect the RNA-binding properties of either of the two proteins and confirms the specificity of our mutations in abolishing the protein-protein interaction between Staufen and Tm1.

Taken together, our data demonstrate that the interaction between Staufen and Tm1 involves hydrophobic residues in Staufen dsRBD4 and Tm1’s disordered N-terminal domain, distinct from those responsible for RNA-binding.

### Interaction of Staufen with Tm1 does not affect their RNA binding

The mutual interaction sites of Tm1 and Staufen are in close proximity to their respective RNA-binding regions (Vaishali *et al*, 2021; Micklem *et al*, 2000; Ramos *et al*, 2000). This raises the possibility that interaction between the proteins could result in cooperative RNA-binding effects, increasing or decreasing in affinity or specificity for *oskar* mRNA. To test this idea, we performed EMSAs with both proteins and the *oskar* region 2+3 RNA. We first titrated increasing concentrations of Tm1 1-335 to *oskar* region 2+3, in presence of a constant concentration of Staufen dsRBD3-4-L4. Under these conditions, *oskar* region 2+3 RNA bound exclusively to Staufen, as no further mobility shifts of the RNA were observed in presence of increasing Tm1 concentrations (Figure 4A). When we titrated increasing amounts of Staufen dsRBD3-4-L4 to *oskar* region 2+3 RNA in the presence of a constant concentration of Tm1 1-335, we observed a mobility shift of the RNA from the Tm1-bound, towards the Staufen-bound species. However, no additional mobility shift corresponding to a ternary complex was observed (Figure 4B), indicating that Staufen displaces Tm1 from the RNA. In summary, we found no evidence for the formation of a ternary complex between *oskar* RNA, Tm1 and Staufen. Instead, Staufen displaced Tm1 from the RNA, indicating a stronger affinity of Staufen for the RNA. These results indicate that Staufen outcompetes Tm1’s interaction with *oskar* RNA rather than cooperatively enhancing its binding.

**Figure 4:**
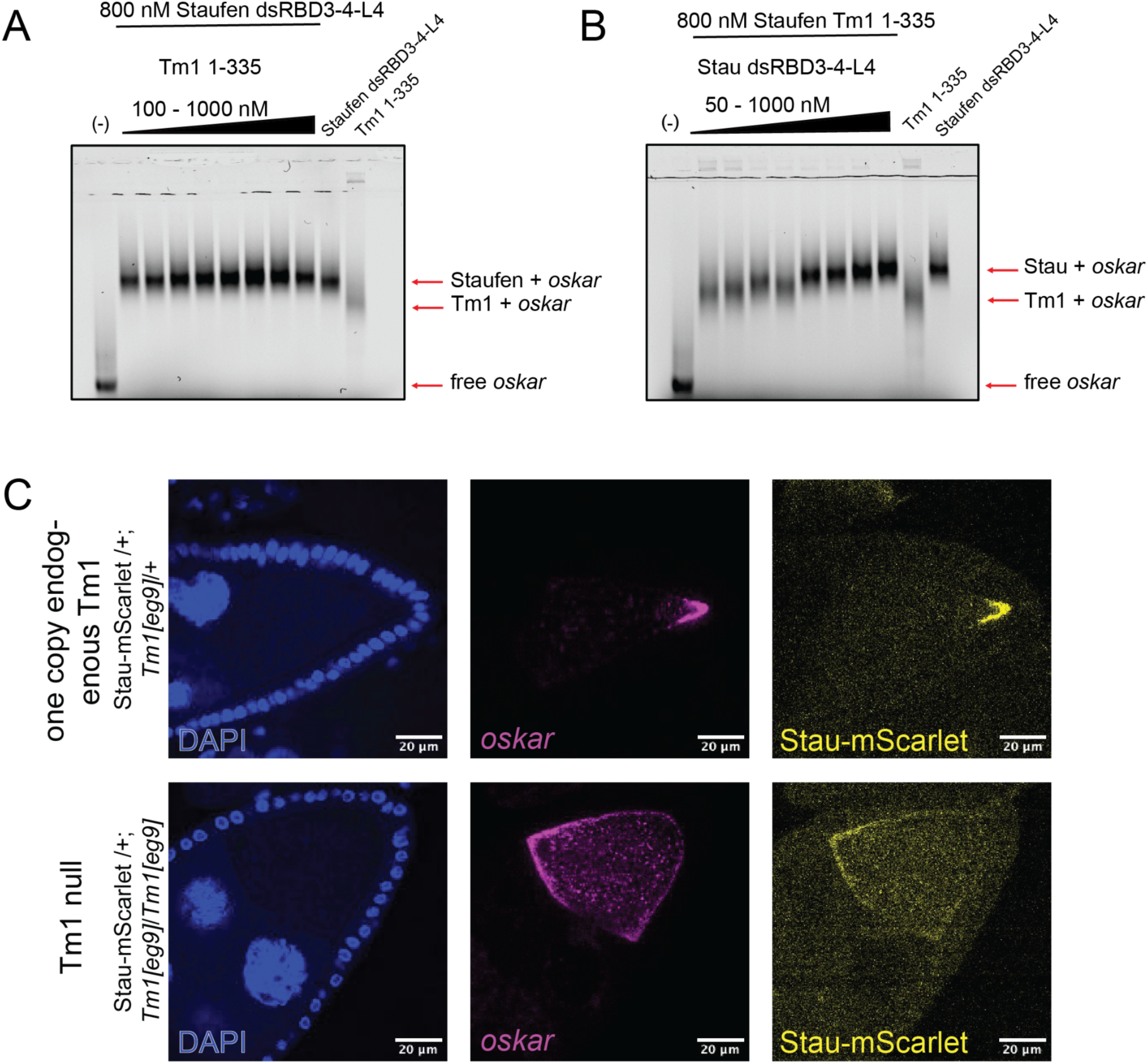
The Staufen-Tm1 interaction does not influence the proteins’ RNA binding. **A**) EMSA with increasing concentrations of Tm1 1-335 (100, 200, 400, 500, 600, 700, 800, 1000 nM) and a constant concentration of Staufen dsRBD3-4-L4 (800 nM); **B)** EMSA with increasing concentrations of Staufen dsRBD3-4-L4 (50, 100, 200, 400, 500, 600, 800, 1000 nM) and a constant concentration of Tm1 1-335 (800 nM); proteins were incubated with *oskar* 3’UTR region 2+3 (Jambor *et al*, 2014; Gáspár *et al*, 2023), which was fluorescently labeled with aminoallyl-UTP-ATTO-680. Free RNA as well as protein/RNA complexes are marked with arrows. **C)** Representative images showing the distribution of *oskar* mRNA (by smFISH) and endogenously tagged Staufen-mScarlet in stage 8/9 *Tm1[eg9]*/+ (upper panel) or *Tm1[eg9]/Tm1[eg9]* (lower panel) oocytes. Nuclei are stained with DAPI. Scale bar: 20 µm.

### Co-localization of Staufen with *oskar* mRNA does not require Tm1

To test the physiological relevance of our *in vitro* findings, we imaged localization of endogenously mScarlet-tagged Staufen in *Tm1[eg9]* mutant *Drosophila* ovaries. In these ovaries, no Tm1 protein is expressed and *oskar* mRNA fails to localize at the posterior of the oocyte, resulting in sterile offspring (Erdélyi *et al*, 1995). We assessed the colocalization of endogenous mScarlet-tagged Staufen with *oskar* mRNA by smFISH in stage 8-9 oocytes. As previously reported (Erdélyi *et al*, 1995), *oskar* localization to the posterior pole was impaired in these flies as compared to flies expressing endogenous Tm1 (Figure 4C). However, Staufen still colocalized with *oskar*, indicating that Staufen recognition of *oskar* mRNA does not require interaction of Staufen with Tm1 (Figure 4C). This finding is consistent with our *in vitro* observation that Staufen outcompetes Tm1 in RNA binding (Figure 4A).

### The Staufen and Tm1 interaction is required for *oskar* mRNA localization and posterior localization of Tm1

Having identified mutations specifically interfering with the interaction between Staufen and Tm1 while leaving the protein structures and their RNA-binding functions intact, we set out to assess the functional significance of the Staufen-Tm1 interaction. To this end, we generated mutations in GFP-tagged Tm1 transgenes and tested them *in vivo*. We expressed the constructs in *Tm1[eg9]* mutant flies, in which endogenous Tm1 is not present and *oskar* mRNA fails to localize (Erdélyi *et al*, 1995; Gáspár *et al*, 2017), and assayed *oskar* mRNA localization by smFISH. As previously shown, wildtype GFP-Tm1 1-334 rescued the grandchildless phenotype and *oskar* localization (Dimitrova-Paternoga *et al*, 2021) (Figure 5A,B,C). Flies expressing the Tm1 M12 mutant in the aa 1-334 context (GFP-Tm1 1-334 M12), produced fertile offspring, but *oskar* localization was partially impaired. While a fraction of *oskar* still enriched at the posterior pole during stage 9 of oogenesis, we often observed an accumulation of *oskar* in a central dot, detached from the posterior cortex (Figure 5A,B). Such a phenotype was previously described as the result of Khc hyperactivity (Williams *et al*, 2014) or an imbalance of Khc and MyoV activities (Krauss *et al*, 2009; Lu *et al*, 2020). Furthermore, we found that colocalization of the transgenic GFP-Tm1 protein with *oskar* RNA at the posterior pole was abolished in the case of the M12 mutant (Figure 5D), indicating that the interaction of Tm1 with Staufen is necessary to retain Tm1 association with *oskar* mRNPs in the oocyte. Of note, the previously described loss of colocalization between *oskar* and an N-terminal deletion mutant of Tm1 (Dimitrova-Paternoga *et al*, 2021), in which the Staufen interaction site is lost, can likely also be explained by the loss of this protein-protein interaction.

**Figure 5:**
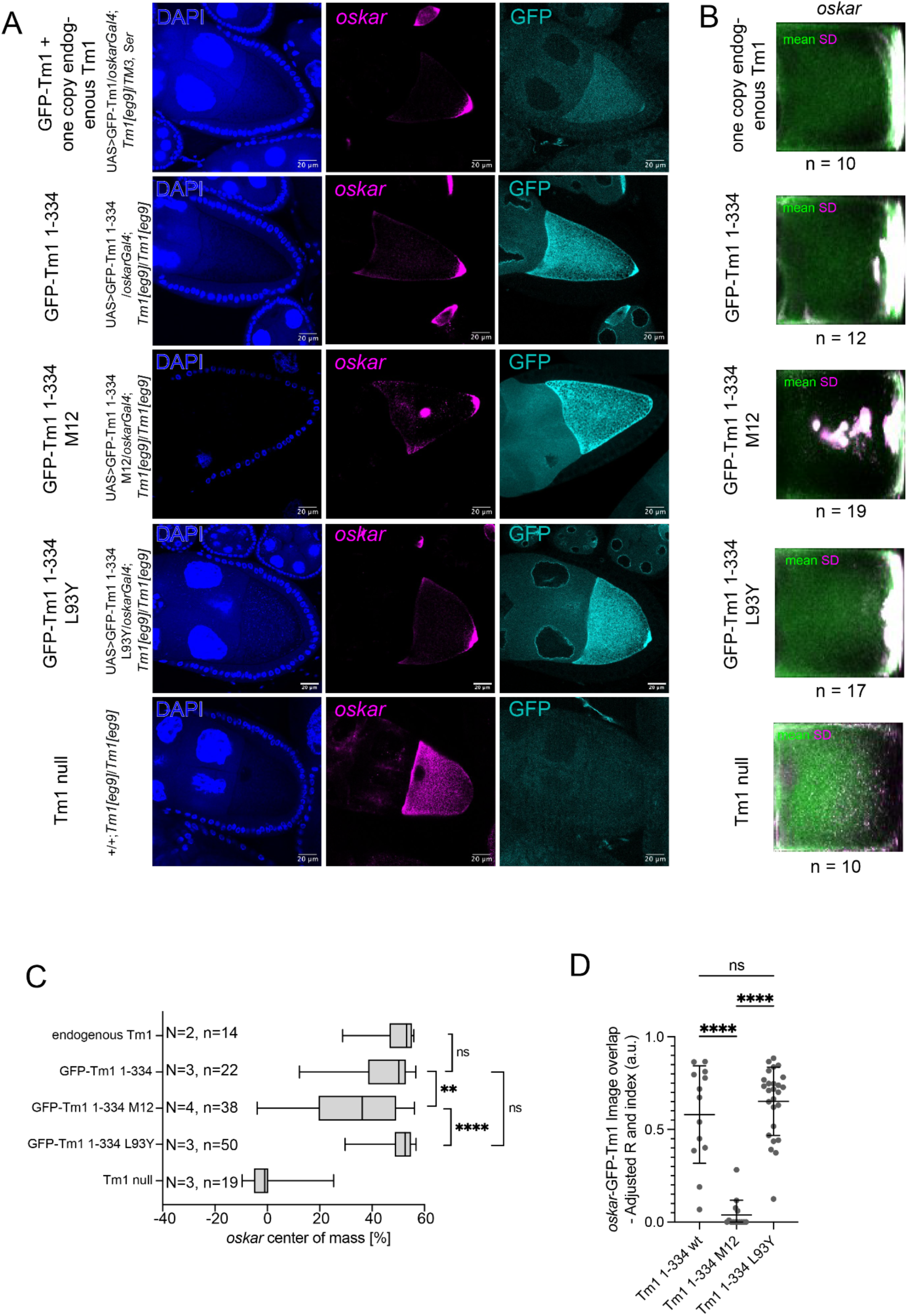
Tm1 deficient of Staufen-binding partially rescues *oskar* mRNA localization but fails to colocalize with *oskar* mRNPs *in vivo*. **A**) Example images showing *oskar* mRNA localization by smFISH and transgenic GFP-Tm1 protein distribution in stage 9 *Tm1[eg9]* oocytes. **B**) Mean *oskar* mRNA distribution and SD within stage 9 *Tm1[eg9]*/*TM3* or *Tm1[eg9]*/*Tm1[eg9]* egg-chambers expressing Tm1 mutant transgenes. **C**) Position of the *oskar* mRNA center of mass relative to the geometric center of the oocyte (0) along the anteroposterior axis. The posterior pole is to the right of the chart. N = number of crosses from which female flies were analyzed; n= number of oocytes analyzed. Bottom and the top of the box represent the first and third quartiles, vertical lines indicate the data median. Whiskers show the entire data range (min to max). *P*-values of pairwise Mann–Whitney *U*-tests are indicated. **D**) Quantification of image overlap for GFP-Tm1 and *oskar* was assessed using *Adjusted Rand index:* A variation of the Rand index, which takes into account the fact that random chance will cause some objects to occupy the same clusters. A score of 1 indicates complete overlap. Statistical significance was assessed by one-way ANOVA followed by Dunnett’s multiple comparisons test. Data points displayed represent single oocytes; error bars indicate mean ± SD. Statistically significant pairwise comparisons are indicated. ****, p <0.0001; **, p <0.01; ns, not significant (p >0.05). Scale bar: 20 µm.

A Tm1 mutant with reduced Staufen-binding, GFP-Tm1 1-334 L93Y, expressed at similar levels (Figure S11), rescued the grandchildless phenotype as well as *oskar* mRNA localization, similar to the wildtype Tm1 transgene (Figure 5A,B,C). Furthermore, this mutant Tm1 variant also colocalized with *oskar* mRNA at the posterior pole (Figure 5D), indicating that a weak interaction between the two proteins is sufficient to tether Tm1 to *oskar* mRNPs.

## Discussion

*oskar* mRNA transport is a paradigm for RNA localization and, with its two-step transport process relying on consecutive actions of dynein and kinesin, for studies of motor protein regulation. However, the mRNP remodeling processes underlying motor switching remain poorly understood. Here, we describe and characterize a previously unknown protein-protein interaction between two regulators of motor protein activity, Staufen and Tm1 in *oskar* mRNPs and test the role of this interaction in *oskar* mRNA localization. In the past, the two proteins have been individually identified as components essential for directed *oskar* mRNA localization (St Johnston *et al*, 1991; Kim-Ha *et al*, 1995; Ephrussi *et al*, 1991; Erdélyi *et al*, 1995; Gáspár *et al*, 2017; Veeranan-Karmegam *et al*, 2016; Dimitrova-Paternoga *et al*, 2021).

Staufen is recruited to the *oskar* mRNP in the oocyte, where the RNP switches from dynein to Khc-mediated transport. Here, we used SPR to show that Staufen is a direct interaction partner of the Khc-regulator Tm1. Using SPR and NMR we mapped the interaction to Staufen dsRBD4 and the disordered N-terminal region of Tm1 (aa 1-213).

Of note, dsRBDs 3 and 4 of Staufen were previously reported to bind RNA (Micklem *et al*, 2000), and RNA-binding mutations in dsRBD3 led to impaired localization of *bicoid* and *oskar* mRNA in the early embryo (Ramos *et al*, 2000). Our selected mutations abolishing the interaction between Staufen and Tm1 do not impact RNA binding by Staufen.

In earlier studies, Tm1 was also shown to display RNA binding (Vaishali *et al*, 2021; Gáspár *et al*, 2017; Dimitrova-Paternoga *et al*, 2021; Sysoev *et al*, 2016). An RNA-binding site in the N-terminus of Tm1 (Vaishali *et al*, 2021) is flanked by residues involved in Staufen-binding (Tm1 aa 20-22 and aa 48-67). Yet, our Staufen binding-deficient Tm1 mutant shows unimpaired RNA binding, indicating that these Tm1 functions can be assessed separately.

Furthermore, despite the spatial proximity of the interaction surfaces in both proteins to their RNA-binding sites, we find no evidence for cooperative RNA binding. Instead, we provide evidence for competitive RNA binding. *In vitro*, Staufen binding to RNA is stronger than Tm1’s binding to RNA, leading to out-competition of Tm1 from *oskar* RNA upon Staufen binding. *In vivo*, Tm1 is displaced from *oskar* mRNPs in the oocyte when its interaction with Staufen is abolished, indicating that its interaction with *oskar* mRNA is weakened under conditions where Staufen is present. Such loss of Tm1 from *oskar* mRNPs was previously described for an N-terminal deletion mutant of Tm1 (Tm1 246-334) (Dimitrova-Paternoga *et al*, 2021). From our study it becomes clear that in this mutant, the interaction site with Staufen is deleted. Hence, the loss of Tm1 from *oskar* mRNPs can also be explained by a loss of this protein-protein interaction.

*oskar* mRNA localization is impaired when Staufen and Tm1 cannot interact and Tm1 is lost from the mRNP. The phenotype we observed for such conditions using our Tm1 M12 mutant resembles that described for Khc hyperactivity or an imbalance between Khc and MyoV activities, pointing to a misregulation of Khc (Williams *et al*, 2014; Krauss *et al*, 2009; Lu *et al*, 2020). Tm1 was previously shown to inhibit Khc activity during dynein-mediated transport via a mechanism involving its N-terminal domain and rearrangement of the Khc stalk domain, stabilizing its autoinhibited conformation (Heber *et al*, 2024). As Tm1 is not displaced from *oskar* mRNPs in the oocyte when Khc is active, it remains unclear how Khc is activated. A likely explanation is a conformational change, possibly by Staufen-induced mRNP remodeling of the Khc-Tm1 complex, promoting Khc activity. Our data also suggest that, upon interaction of the Tm1 N-terminal domain with Staufen in the oocyte, the regulatory interaction between the Khc tail and Tm1’s N-terminal domain (Heber *et al*, 2024) is prevented, thereby allowing Khc to switch to its active conformation (Figure 6). Tm1’s interactions with Khc and possibly *oskar* mRNA are thus weakened and Tm1 is retained in the mRNP through its interaction with Staufen, in close proximity to Khc. As loss of Tm1 impairs *oskar* localization in a pattern resembling that of hyperactive Khc mutants, its retention in the mRNP may allow fine-tuning of Khc activity in the oocyte by Tm1-promoted interactions within the Khc stalk (Heber *et al*, 2024). As Staufen was previously shown to inhibit dynein by displacing Egl from the RNP when motor switching becomes necessary (Gáspár *et al*, 2023), an additional role of Staufen in Khc activation would constitute an accurate and efficient way to couple dynein and kinesin motor regulation during the switch. It will be interesting to understand how Staufen-induced changes of the mRNP are linked to other factors required for Khc-mediated transport, such as the EJC-SOLE complex (Ghosh *et al*, 2012) or Ensconsin (Sung *et al*, 2008).

**Figure 6:**
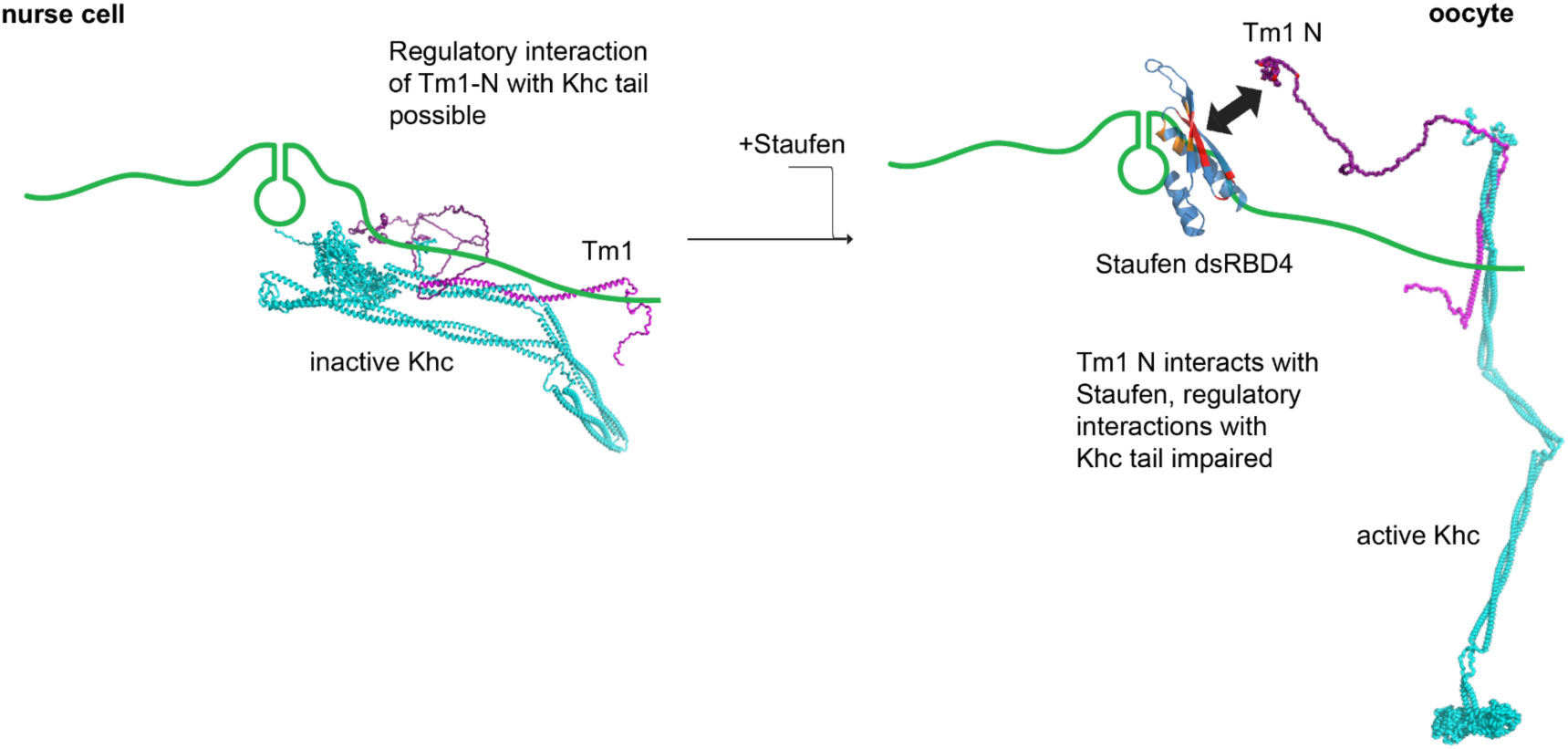
Model of the interactions of Staufen and Tm1 in *oskar* mRNP transport. During dynein-mediated transport in the nurse cell, Tm1 keeps Khc in an inactive conformation. Tm1 achieves this via its N-terminal domain and Khc’s regulatory tail (Heber *et al*, 2024), and links Khc to the mRNA via an interaction within both proteins’ coiled-coil domains (Dimitrova-Paternoga *et al*, 2021). Possibly also an interaction between the Tm1 N-terminal domain and the RNA (Vaishali *et al*, 2021) contributes to Tm1’s function as a Khc adaptor. In the oocyte, Staufen binds to the *oskar* mRNA and the mRNP is remodeled to allow for Khc activity. The N-terminal region of Tm1 interacts with Staufen dsRBD4, which prevents it from regulating Khc. As Tm1’s N-terminal interactions with Khc and RNA are thus prevented, the additional interaction with Staufen is necessary to retain Tm1 associated with the mRNP and thus in close proximity to Khc. In summary, these interactions allow for the fine-tuned activation of Khc. Red: residues involved in the Tm1-Staufen interaction. Orange: residues involved in RNA-binding.

Our Staufen-binding deficient Tm1 transgene rescued the grandchildless *Tm1*[*eg9]/Tm1[eg9]* phenotype (Erdélyi *et al*, 1995), indicating that *oskar* localization is rescued to an extent sufficient for pole cell formation. As *oskar* localization in the female germline is crucial for viability and fertility of the fruit fly, it is likely that several redundant mechanisms ensure the mRNA’s enrichment at the posterior pole. The interaction between Tm1 and Staufen might be one of many weak or transient interactions that cooperatively ensure assembly and remodeling of the *oskar* mRNP. Likely, such redundant, individual interactions are important to ensure correct RNA localization also under suboptimal conditions such as lack of nutrients or temperature stress, to ensure progression through oogenesis and correct development of the resulting embryo.

## Materials & Methods

Key resources table.

Key resources table
| Reagent/Resource | Source | Identifier |
| --- | --- | --- |
| **Plasmids** |  |  |
| pBluescript-Blue-*oskar* | Anne Ephrussi | N/A |
| pFastBacDual-Staufen FL | This work | N/A |
| pGEX-6P-1-Staufen Nterm-dsRBD1-2 (aa 1-557) | This work | N/A |
| pGEX-6P-1-Staufen dsRBD3-4-5-Cterm (aa 578-1026) | This work | N/A |
| pGEX-6P-1-StaufendsRBD3-4-L4 (aa 578-950) | This work | N/A |
| pGEX-6P-1-Staufen dsRBD3-4 (aa 578-785) | This work |  |
| pGEX-6P-1-Staufen L4-dsRBD5-Cterm (aa 782-1026) | This work | N/A |
| pGEX-6P-1-Staufen dsRBD3 (aa 578-645) | This work | N/A |
| pGEX-6P-1-Staufen dsRBD4 (aa 711-781) | This work | N/A |
| pGEX-6P-1-Staufen L4-dsRBD5 (aa 951-1026) | This work | N/A |
| pGEX-6P-1-Staufen dsRBD3-4-L4-M6 (RTY-FSV) | This work | N/A |
| pETM11-Tm1 FL | Dimitrova-Paternoga *et al*, 2021 |  |
| pGEX-6P-1 Tm1 FL | This work | N/A |
| pETM11-Tm1 1-335 | Dimitrova-Paternoga *et al*, 2021 | N/A |
| pGEX-6P-1 Tm1 1-335 | This work | N/A |
| pETM11-Tm1 1-213 | Dimitrova-Paternoga *et al*, 2021 | N/A |
| pGEX-6P-1-Tm1 1-213 | This work | N/A |
| pGEX-6P-1-Tm1 1-213 M12 | This work | N/A |
| pGEX-6P-1-Tm1 1-213 M3 | This work | N/A |
| pGEX-6P-1-Tm1 1-213 M8 | This work | N/A |
| pGEX-6P-1-Tm1 1-213 L93Y | This work | N/A |
| pGEX-6P-1-Tm1 1-335 M12 | This work | N/A |
| pFastBacDual-HisSUMO-SNAP-3C-Khc | Heber *et al*, 2024 | N/A |
| pFastBacDual-HisSUMO-SNAP-3C-Khc-Tm1 | Heber *et al*, 2024 | N/A |
| pET-Duet 1-His-SNAPf-Btz1-345-eIF4AIII | This work | N/A |
| pET-Duet 1-His-Mago-SNAPf-Y14 | This work | N/A |
| pUASpattB-emGFP-Tm1 1-334 M12 | This work | N/A |
| pUASpattB-emGFP-Tm1 1-334 L93Y | This work | N/A |
| **Proteins** |  |  |
| 3C-protease | In house | N/A |
| **Fly strains** |  |  |
| *Tm1[eg9]* (FBal0049223) | Erdélyi *et al*. 1995 | N/A |
| *oskarGAL4/TM3, Sb* | BDSC | BDSC Cat# 44242; RRID:BDSC_51324 |
| emGFP-Tm1 1-334/CyO | Dimitrova-Paternoga et al., 2021 | N/A |
| emGFP-Tm1 1-334 M12/CyO | This study | N/A |
| emGFP-Tm1 1-334 L93Y/CyO | This study | N/A |
| Staufen::mScarlet | Gift of Akira Nakamura | N/A |

### Molecular Cloning

For GST-fusion constructs, empty pGEX-6P-1 plasmids were linearized using *Bam*HI and *Xho*I (NEB, MA, USA) and PCR-amplified inserts ligated according to standard protocols. For seamless cloning, pFastBacDual vectors were linearized with *Bam*HI and *Hind*III (NEB, MA, USA) and inserts ligated using the InFusion® HD Cloning Kit (Takara, Japan), according to manufacturer’s instructions. Inserts were amplified by PCR using Q5® High-Fidelity DNA Polymerase (NEB, MA, USA)

Core components of the EJC were cloned into the pET-Duet vector for coexpression of eIF4AIII-Btz 1-345 and Y14-Mago heterodimers, respectively. The respective coding sequences were subsequently cloned into multiple cloning sites 1 between the *Bam*HI and *Not*I sites and 2 between *Nde*I and *Xho*I sites of the linearized vector.

GFP-Tm1 1-334 mutant vectors for fly transgenesis were made by site-directed mutagenesis of pUASpattB-emGFP-Tm1 1-334 (Dimitrova-Paternoga et al., 2021). All plasmids were verified by Sanger sequencing (Eurofins, Germany).

### Protein expression and purification

#### Protein expression in insect cells

Recombinant full-length, GST-tagged Staufen was expressed and purified from insect cells (H5) using the baculovirus expression system to yield close-to-physiological state proteins. The plasmid pFastBacDual-Staufen FL was introduced into DH10bac *E. coli* chemically competent cells (Inoue *et al*, 1990), followed by blue-white screening for correct insertion of the construct of interest into the bacmid. Obtained bacmids were isolated and verified by Sanger sequencing (Eurofins, Germany). To produce the recombinant virus, Sf21 cells were transfected with bacmid in 6-well plates containing 0.4×10^6^ cells/mL. The transfection was carried out using FuGENE® HD Transfection Reagent (Promega) as per the manufacturer’s instructions. After 96h, P_0_ was harvested from the media supernatant. P_1_ was obtained by infecting 10 mL Sf21 cells with 4 mL P_0_ and harvesting by centrifugation at 2,000 g for 10 min after 72 h. Similarly, P_2_ was obtained by infecting 250 mL Sf21 cells with the complete volume of P_1_ for 96 h and harvesting the supernatant by centrifugation at 2,000 g for 10 min and filtration (0.2 µm). To produce the GST-tagged Staufen with P_2_, 40 mL of the virus was used to infect 1,000 mL of H5 cells at 1×10^6^ cells/mL in a shaking culture with 80 rpm at 27.5 °C. After 72 h, cells were harvested by centrifugation at 2,000 g for 10 min. The pellet was resuspended in Lysis Buffer (1 M NaCl, 20 mM Tris, 5% Glycerol, 0.01% Tween-20, 2 mM MgCl₂, 2 mM DTT), flash-frozen in liquid nitrogen, and stored at –80 °C.

Khc and Khc-Tm1 full-length proteins were expressed using the Bac-to-Bac Baculovirus expression system (Gibco) in Sf21 cells as described before (Heber *et al*, 2024).

#### Protein expression in *E. coli*

Plasmids were transformed into *E. coli* BL21 Gold (DE3) and plated on 2-YT agar plates containing ampicillin (100 µg/mL) or kanamycin (50 µg/mL). For liquid culture, a single colony was selected after overnight incubation at 37 °C. *E. coli* was grown until O.D.600 of 0.8. Protein expression was induced by 0.5 mM IPTG. After induction, the cultures were cooled to 18 °C and cultivated overnight. The cultures were harvested by centrifugation 24 h post-induction, flash-frozen in liquid nitrogen, and stored at –80 °C.

^15^N– and ^15^N-^13^C-labeled proteins were expressed in *E. coli* BL21 (DE3) cells in M9 media after induction with IPTG at 18 °C for 16 h.

#### Purification of Staufen proteins

*E. coli* cells expressing recombinant Staufen proteins and mutations were suspended in lysis buffer (1 M NaCl, 20 mM HEPES pH 7.5, 2 mM MgCl_2_, 2 mM DTT) supplemented with 1x tablet of Complete Mini Protease Inhibitor cocktail (Roche, Germany), 5 % glycerol and 0.1 % Tween-20. Lysis was accomplished using an LM10 Microfluidizer (Microfluidics, MA, USA). Lysate was centrifuged at 30,000 *g* at 4 °C for 20 min to remove insoluble components and cell debris. The supernatant was clarified using a 0.45 µm filter, and proteins with a GST tag were affinity purified using a 5 mL GSTrap FF (Cytiva, MA, USA). After injection of the clarified supernatant, bound proteins were washed with lysis buffer until baseline was reached.

Cleavage of the GST-tag was accomplished on-column by injecting 5 mL 3C protease (0.2 mg/mL) in lysis buffer followed by incubation at 4 °C overnight. The cleaved protein was eluted in lysis buffer. To avoid any leakage of cleaved GST-tag and 3C protease, the eluate was passed over a second 5 mL GSTrap FF column. Eluted fractions were analyzed by SDS-PAGE and fractions containing the protein of interest were dialyzed against dialysis buffer (150 mM NaCl, 20 mM HEPES pH 7.5, 2 mM DTT) at 4 °C overnight, using a 3.5 kDa MWCO membrane (Roth, Germany) to reduce the NaCl concentration. After dialysis the sample was applied to a HiTrap Heparin HP column (Cytiva, MA, USA) equilibrated in the dialysis buffer. Elution was accomplished using a 15 CV gradient from 0-100 % heparin elution buffer (1 M NaCl, 20 mM HEPES pH 7.5, 2 mM DTT). Fractions were analyzed by SDS-PAGE and desired fractions were subjected to size exclusion chromatography using a HiLoad 16/600 Superdex 200 pg column (Cytiva, MA, USA) equilibrated in the dialysis buffer or on a HiLoad 16/600 Superdex 75 pg column (Cytiva, MA, USA) in NMR buffer (25 mM NaPO4 pH 6.5, 150 mM NaCl and 0.5 mM Tris(2-carboxyethyl)phosphine for NMR experiments. Fractions were analyzed by SDS-PAGE, pooled, and concentrated using an Amicon® 10 kDa MWCO filter (Merck, Germany). Protein concentrations were measured at 595 nm using Bradford assay or at 562 nm by BCA assay. 50 µL protein aliquots were flash-frozen in liquid nitrogen and stored at –80 °C.

For the purifications of Staufen dsRBD3 and Staufen dsRBD4 the Heparin column was omitted.

#### Purification of Tm1 proteins

Recombinant Tm1 proteins and mutations expressed in *E. coli* were purified essentially as described previously (Heber *et al*, 2024; Dimitrova-Paternoga *et al*, 2021). After harvest, cells were suspended in lysis buffer (500 mM NaCl, 20 mM HEPES pH 7.5, 2 mM DTT) and supplemented with 1 tablet of Complete Mini Protease Inhibitor cocktail (Roche, Germany). The lysis, affinity chromatography, and size exclusion chromatography were accomplished as described previously (Heber *et al*, 2024; Dimitrova-Paternoga *et al*, 2021). Fractions were analyzed by SDS-PAGE and desired fractions were pooled and concentrated using an Amicon® 10 kDa MWCO filter (Merck, Germany). Protein concentrations were measured at 595 nm using Bradford assay. 50 µL protein aliquots were flash-frozen in liquid nitrogen and stored at –80 °C.

#### Purification of Khc and Khc-Tm1 full-length proteins

Purification of Khc and Khc-Tm1 full-length proteins was performed as described before (Heber *et al*, 2024). Briefly, after expression in Sf21 cells for 3 days at 27.5 °C, cells were harvested by centrifugation at 1,000 *g* for 20 min at 4 °C. The pellets were lysed in a dounce tissue grinder in lysis buffer (20 mM Tris–HCl at pH 7.5, 500 mM NaCl, 1 mM MgCl_2_, 0.1 mM ATP, 2 mM DTT, 5% glycerol, 0.2% Tween-20). The lysates were cleared by centrifugation and the soluble protein fraction was affinity-purified on a HisTrap Excel column (GE Healthcare). After elution in a 0–300 mM imidazole gradient, the HisSUMO-fusion tag was cleaved by 3C protease during dialysis in GF150 buffer (25 mM HEPES/KOH pH 7.3, 150 mM KCl, 1 mM MgCl2, 0.1 mM ATP, and 2 mM DTT) for 16 h before further purification by anion-exchange chromatography on a HiTrap Q HP column (GE Healthcare), followed by size-exclusion chromatography (SEC) on a Superdex 200 Increase 10/300 column (GE Healthcare) in GF150 buffer.

#### Purification of EJC components

*Drosophila* EJC components were expressed and purified essentially as described before for the human EJC (Bono *et al*, 2006) Briefly, Btz-SELOR/eIF4AIII and Mago/Y14 were co-expressed from a bicistronic vector as His-tagged fusion proteins and co-purified by Ni-IMAC on HisTrap FF (Cytiva) followed by tag-cleavage using 3C-protease and size exclusion chromatography on a HiLoad Superdex 200 16/600 column in GF150 buffer.

## Surface plasmon resonance

### Antibody coupling

CM5 sensor chips (Cytiva, MA, USA) were coupled with anti-GST antibody (Cytiva, MA, USA) using the GST Capture kit (Cytiva, MA, USA) according to the manufacturer’s protocol included in the Biacore S200 software. Both flow cells and reference cells were coupled with anti-GST antibody (Cytiva, MA, USA).

### Amine coupling

The ligands were directly coupled to CM5 sensor chips by utilizing the Amine Coupling kit (Cytiva, MA, USA) according to the manufacturer’s protocol included in the Biacore S200 Control software (Version 1.1). Prior to the amine coupling, pH scouting was done for each protein in order to determine optimal coupling conditions based on the manufacturer’s protocol. Blank coupling was performed on the reference cell.

### Single-cycle kinetics

Before injecting the analyte, GST was captured by the GST antibody on the reference cell, and the corresponding GST-fusion protein on the flow cell. The analytes were diluted in HBS EP buffer (150 mM NaCl, 10 mM HEPES pH 7.4, 3 mM EDTA, 0.05 % Tween-20). 0.1 % SDS in H_2_O was used for regeneration. The pumps were primed before the first injection, and a startup cycle containing only HBS EP was performed. The analysis temperature was set to 25 °C, and the sample compartment was cooled to 12 °C. The flow rate was set to 30 µL/min at a data collection rate of 10 Hz. Reference cell subtracted data were used to determine the equilibrium dissociation constants (K_D_) in the Biacore Evaluation software (Version 1.1) performing a one-site binding fit of at least six individual concentrations. Every single-cycle kinetic experiment that indicated an interaction between the two tested proteins was repeated twice. The K_D_ value is presented as the mean with the standard deviation indicated by ±.

### Multi-cycle kinetics

The analytes were diluted in HBS EP buffer supplemented with 150 mM NaCl. For regeneration, a solution of 0.1 % SDS in H_2_O was used. Prior to the first injection, the pumps were primed and a startup cycle with HBS EP was performed. The sample compartment was cooled to 12 °C, and the analysis temperature was set to 25 °C. The flow rate was set to 30 µL/min and the experiment was conducted at a data collection rate of 10 Hz. The final results were obtained after reference cell subtraction. K_D_ values were calculated using the Biacore Evaluation software (Version 1.1) using a one-site binding fit based on at least eleven individual concentrations. Double measurements for one concentration were included in each run, and each multi-cycle kinetic experiment was repeated twice. K_D_ values are shown as the mean of all three replicates, with ± indicating standard deviation.

## NMR spectroscopy

NMR measurements were performed at 298 K on a Bruker Avance III NMR spectrometer with magnetic field strengths corresponding to proton Larmor frequencies of 700 MHz equipped with room temperature HCN probe head with a z-axis pulsed field gradient. NMR sample concentration was 50 µM for titration experiments and 200 µM for backbone assignment experiments. Experiments for backbone assignments were performed on ^13^C,^15^N-labeled samples using conventional triple-resonance experiments (HNCO, HNCA, CBCA(CO)NH, HN(CO)CA, and HNCACB) (Simon *et al*, 2010; Sattler, 1999). All spectra were acquired using the apodization weighted sampling scheme (Simon & Köstler, 2019), processed using NMRPipe (Delaglio *et al*, 1995) and analyzed using NMRView (Johnson & Blevins, 1994). As the resonances corresponding to the flexible linker L4 dominated the ^1^H-^15^N HSQC spectrum of Staufen dsRBD3-4-L4, masking resonances from the folded dsRBDs, we used a ^15^N-^13^C-labeled Staufen construct comprising dsRBDs 3 and 4 (Staufen dsRBD3-4) for backbone assignment. However, in this construct many dsRBD4 resonances showed severe line-broadening and were missing in the backbone assignment experiments and we could only assign a small subfraction of residues in dsRBD4. A comparison of the ^1^H-^15^N-HSQC spectra of Staufen dsRBD3-4 and and dsRBD3-4-L4 showed that the folded domains remained unaffected by truncation of the linker and assignments based on Staufen dsRBD3-4 can be transferred to the longer protein construct (Figure S12).

For titrations, interaction partners were added to ^15^N-labeled proteins at indicated ratios and a ^1^H,^15^N HSQC was recorded for each titration point. Peak intensity ratios were derived using NMRView (Johnson & Blevins, 1994) and corrected for dilution. The extent of amide ^1^H-^15^N chemical shift perturbations (CSPs) in free versus bound proteins was calculated according to Williamson (Williamson, 2013) to compensate for frequency range differences between ^15^N and ^1^H dimensions.

Errors of individual measurements are estimated as the standard deviation of the noise in a spectral region without any signals. Error bars of intensity ratios are calculated according to the rule of propagation of errors.

## RNA *in vitro* transcription and Electrophoretic Mobility Shift Assay (EMSA)

EMSA was carried out as described previously (Bose *et al*, 2022). In brief, RNA was transcribed *in vitro* using the HiScribe® T7 High Yield RNA Synthesis Kit (NEB, MA, USA) according to the manufacturer’s instructions. Templates for *in vitro* transcription were produced by PCR from pBluescript-*oskar* (Primers: T7-a *oskar* 3’UTR FW: AATTTAATACGACTCACTATAGGGTTGGGTTCTTAATCAAGATAC; T7-*oskar* region 2+3 FW: AATTTAATACGACTCACTATAGGTGTTCTATATACTTTTGTGTGGGTCA; *oskar* 3’UTR RV: ACAAAAAAAACGTGATCACC). Fluorescent labeling was accomplished by adding 1 mM aminoallyl-UTP-ATTO-680 (Jena Bioscience, Germany) to the reaction. The template was digested by TURBO^TM^ DNase (Thermo Fisher Scientific, MA, USA). RNA was phenol-chloroform-isoamyl alcohol extracted, precipitated using 1/10 volume of 3 M sodium acetate and 3 volumes of ice-cold ethanol and dissolved in water. For EMSAs, 50 nM labeled *oskar* RNA were added to each sample with increasing protein concentration. Complexes were allowed to form for 20 min at room temperature in EMSA buffer (150 mM NaCl, 20 mM HEPES pH 7.5, 5 % glycerol). Reactions were resolved in 0.8 % agarose gels at constant 100 V at 4 °C. Fluorescent gel imaging was performed in a LI-COR Odyssey DLx (LI-COR Biosciences, NE, USA).

## Fly genetics and husbandry

Fly experiments were performed essentially as described in Dimitrova-Paternoga *et al*, 2021. Tm1 transgenic flies were generated by site-specific integration of the respective pUASp-attB plasmid (see above) with Φc31 integrase in VK18 {vas-phi-ZH2A, PBac(y[+]-attP-9A)VK00018, Bellen laboratory} fly line for site-specific insertion into the *attP* landing site on chromosome II. The transgenes were balanced with *CyO*. emGFP-Tm1 transgenes were driven by one copy of *oskar*-*Gal4* ((Telley *et al*, 2012); FBtp0083699) in the *Tm1[eg9]*/*Tm1[eg9]* background (FBal0049223) (Erdélyi *et al*, 1995). All fly stocks were grown at 21°C–25°C in vials on standard cornmeal agar. Prior to dissection, freshly one to five days old female flies were fed with dried yeast for one day.

## Imaging experiments

### Single-molecule fluorescent *in situ* hybridization (smFISH)

Forty-two probes against the *oskar* mRNA coding region and 3′ UTR were labeled with Atto633 according to Gaspar *et al*, 2017. smFISH was performed essentially as described in Gáspár *et al*, 2017. Pairs of *Drosophila* ovaries were fixed with 2% (v/v) PFA and in PBS (pH 7.4) for 15 min on an orbital shaker, followed by three washes with PBT (PBS + 0.1% [v/v] Triton X-100 at pH 7.4) for 5 min each. Ovaries were then prehybridized in 200 μL of hybridization buffer (300 mM NaCl, 30 mM sodium citrate at pH 7.0, 15% [v/v] ethylene carbonate, 1 mM EDTA, 50 μg/mL heparin, 100 μg/mL salmon sperm DNA, 1% [v/v] Triton X-100) for 5 min at 37 °C, before 50 μL of probe mixture (25 nM per individual oligonucleotide in hybridization buffer) was added for an additional 3 h at 37 °C. After hybridization, excess probes were removed by two washes in the hybridization buffer (15 min at 37 °C each) and three washes in PBT (5 min at RT each). The samples were then mounted in ProLong™ Gold Antifade Mountant (Invitrogen).

### Microscopy and image analysis

Images were collected on a Leica Stellaris-8 confocal microscope on an inverted DMI8 stand controlled by the LAS X (Leica) software with a Plan Apo 63x/1.40 oil immersion objective (Leica). The microscope was equipped with a white light laser (Leica) that was tuned to the respective excitation wavelengths of each fluorophore. GFP-tagged Tm1 variants were detected via their native GFP-fluorescence, endogenously mScarlet-tagged Staufen was detected via its native mScarlet-fluorescence. The pinhole was set to 1 Airy Units.

Image analysis was performed in ImageJ/Fiji. Analysis of the *oskar* mRNA center of mass distribution was performed according to Gaspar *et al*, 2014. For *oskar*-Tm1 colocalization, images were analyzed using the boutique image analysis software CellProfiler developed by the BROAD institute (V4.2.6, https://cellprofiler.org/). Modules used and optimized for each individual experiment included: Identify primary objects, identify secondary objects, threshold and calculate measure image overlap. Plots and statistical analysis were done with GraphPad Prism 8.

## Competing interest statement

The authors declare no competing interests.

## Data availability

The backbone chemical shift assignments of Staufen dsRBD3-4 were deposited to the BMRB under the accession code 52764. Source data are provided with this paper. All other data is available upon request.

## Acknowledgments

We thank L. Dimitrova-Paternoga (European Molecular Biology Laboratory (EMBL) Heidelberg) and Akira Nakamura (Kumamoto University, Kumamoto, Japan) for reagents, the EMBL Protein Expression and Purification Core Facility and the FMI Structural Biology Platform, as well as the FMI Facility for Advanced Imaging and Microscopy and J. Felsenberg (FMI Basel) for the use of his fly laboratory.

S.H. was supported by the EMBL Interdisciplinary Postdoctoral fellowship (EIPOD) Programme under Marie Curie Cofund Actions MSCA-COFUND-FP (grant no. 664726) and Deutsche Forschungsgemeinschaft (DFG)-Forschergruppe 2333 grant to A.E (grant no. EP37/4-1) and to D.N. (grant no. NI1110/6-1 and 2). Work in the laboratories of J.H. and A.E. was supported by funding from the DFG via the priority program SPP1935 to J.H. and A.E. (grant nos EP37/3-1 and EP37/3-2) and the EMBL.

## Author contributions

S.H., A.E. and D.N. conceived the study. T.G., S.H., J.G., B.S., T.M. and V.R. designed and performed the experiments. T.G., S.H., T.W., B.S., T.M. and J.H. analyzed the data. S.H., J.C., D.N. and A.E. supervised the study. J.C., J.H., D.N. and A.E. acquired funding. T.G. and S.H. drafted the manuscript. All authors contributed to the writing of the manuscript.

## Notes

### Competing Interest Statement

The authors have declared no competing interest.

