## Supplementary Materials for "A direct interaction between the RNA-binding proteins Staufen and Tm1-I/C regulates *oskar* mRNP composition and transport"

### Supplementary data

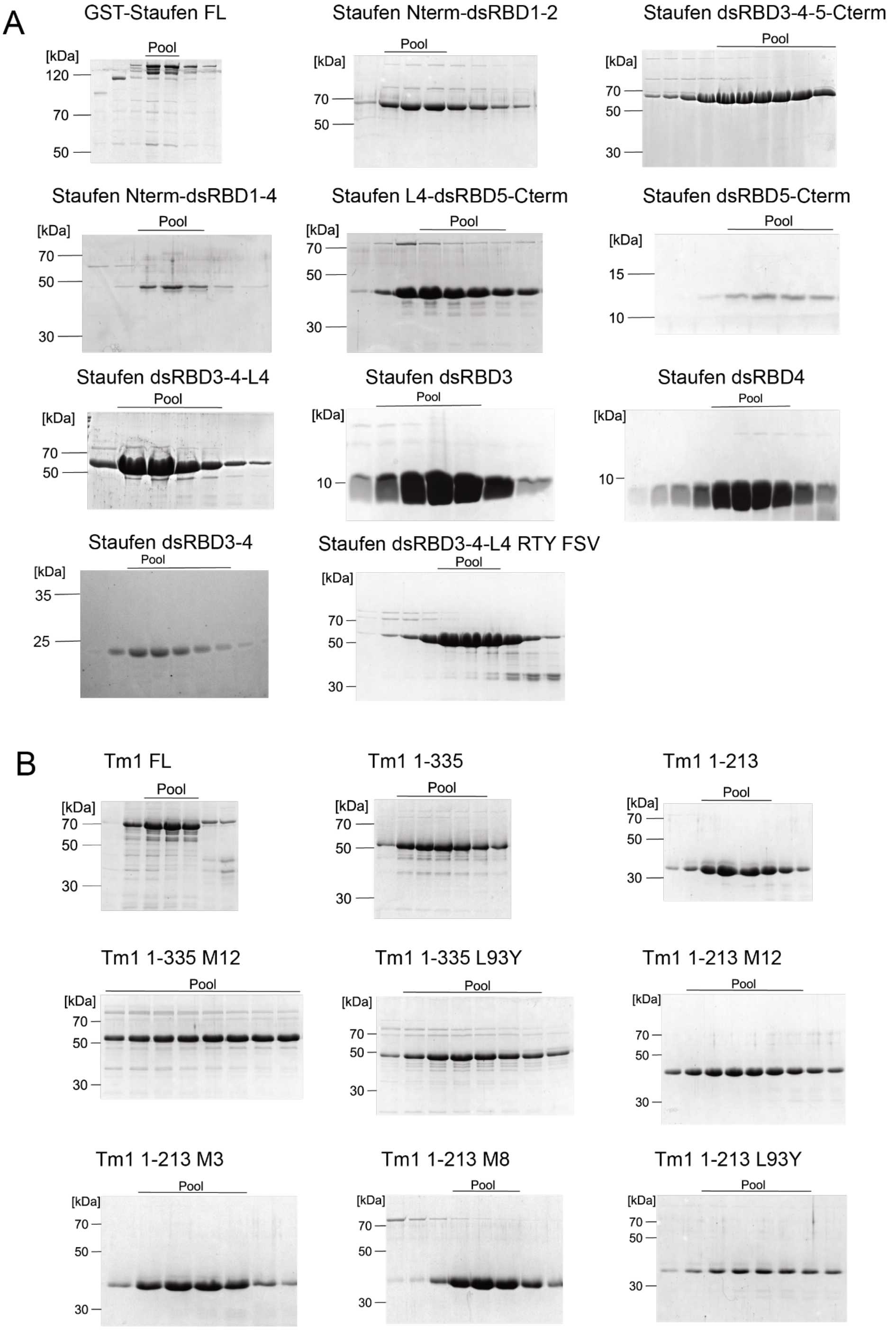


**Figure S1: Purified Staufen and Tm1 proteins used in this study. A)** Exemplary SDS-PAGE of final size exclusion chromatography purification step for all used Staufen proteins and **B)** for all used Tm1 proteins. Fractions pooled for subsequent experiments are indicated.

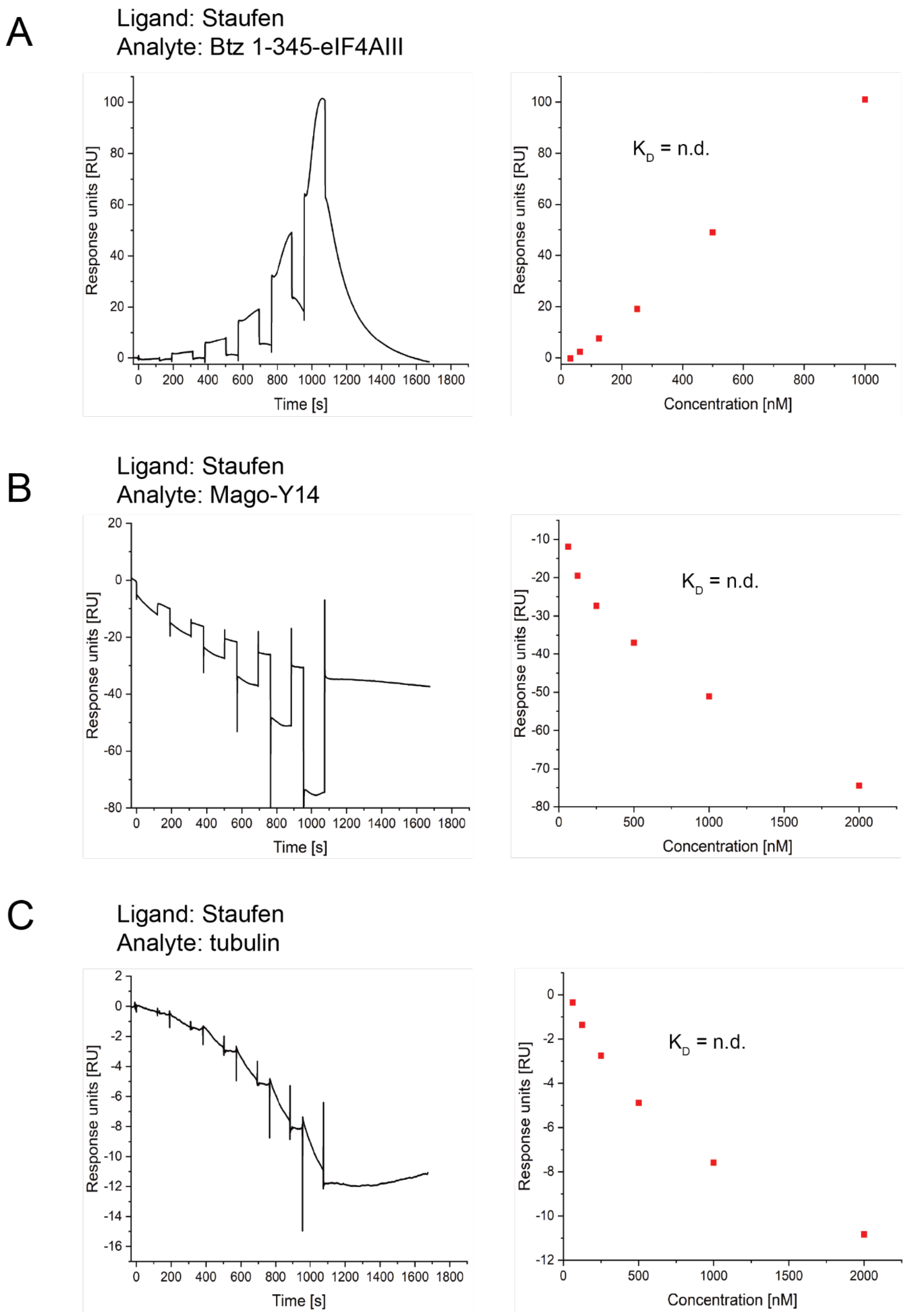


**Figure S2: In SPR no interaction between Staufen and EJC components or tubulin are detected. A-C)** Single-cycle experiment sensorgrams and response concentration plot for Btz SELOR-eIF4AIII (A), Mago-Y14 (B) and tubulin (C) injected onto surface coupled Staufen FL. Due to a lack of concentration-dependent saturation, no equilibrium dissociation constants ($K_D$) could be determined.

A Ligand: Tm1 FL
Analyte: Staufen dsRBD1-2

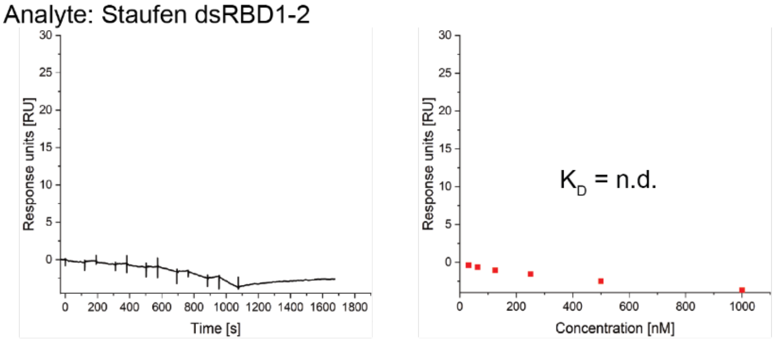


B Ligand: Tm1 FL
Analyte: Staufen dsRBD3-4-5-C-term

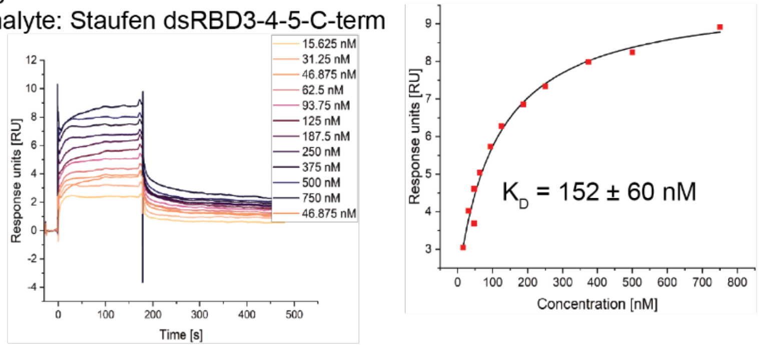


C Ligand: Tm1 FL
Analyte: Staufen dsRBD1-4

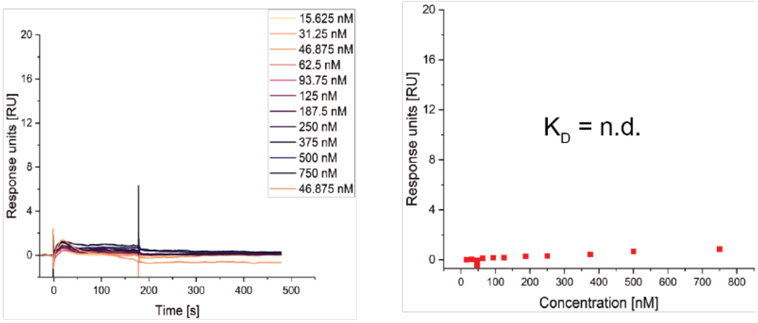


D Ligand: Tm1 FL
Analyte: Staufen dsRBD5-Cterm

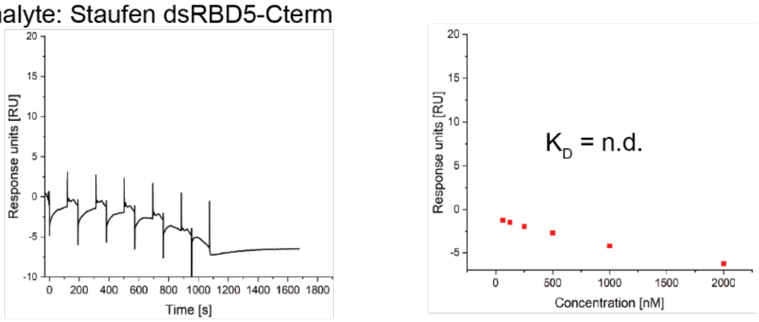


E Ligand: Tm1 FL
Analyte: Staufen L4-dsRBD5-Cterm

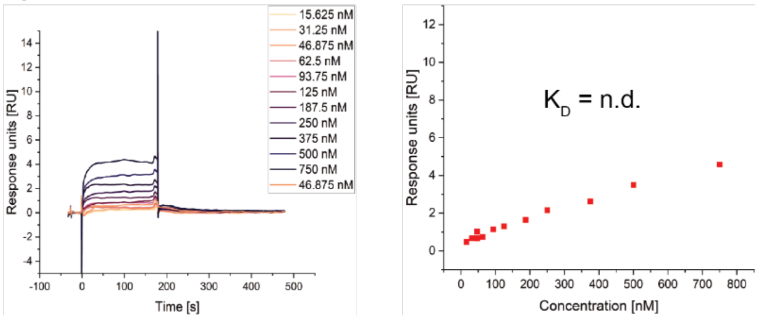

**Figure S3: Staufen fragments showing no binding to Tm1 FL. A-E)** Sensorgrams (left) and response concentration plots (right) of SPR experiments with Staufen dsRBD1-2 (A), Staufen dsRBD3-4-5-C-term (B), Staufen dsRBD1-4 (C), Staufen dsRBD5-Cterm (D) and Staufen L4-dsRBD5-Cterm (E) show no concentration-dependent saturation of binding to surface-coupled Tm1 FL. $K_D$s for Staufen dsRBD3-4-5-C-term was determined using a one-site-binding fit to the maximal response at equilibrium for each analyte concentration. The $K_D$-value is given as mean ± SD from triplicate experiments. For other Staufen proteins, $K_D$s could not be determined.

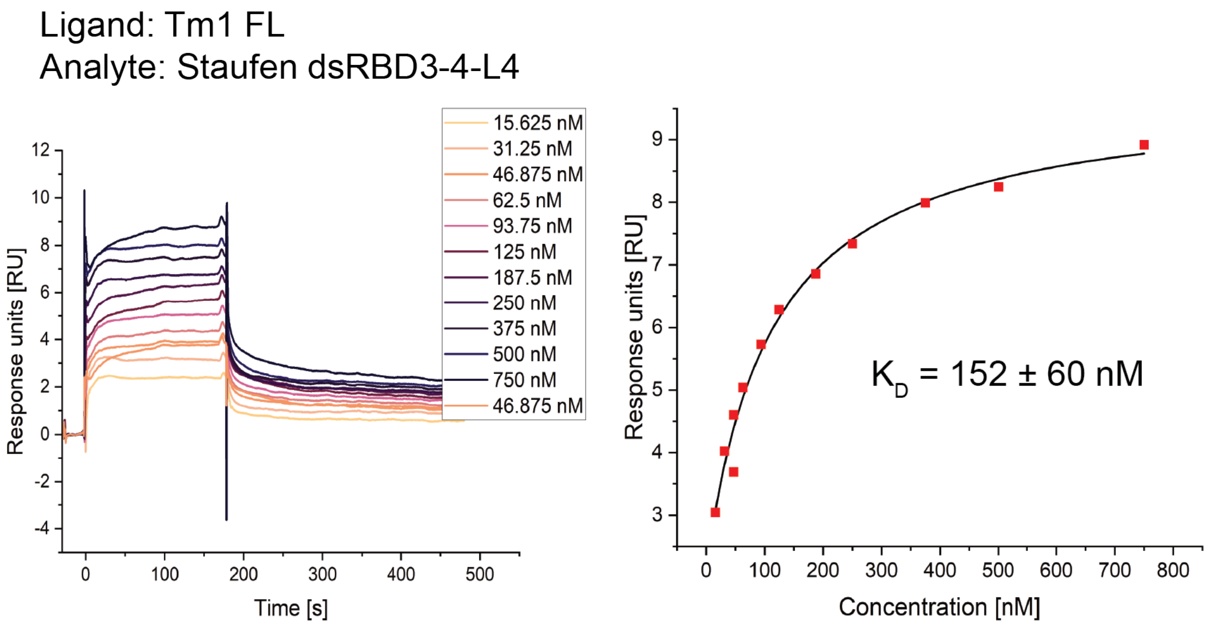


**Figure S4: In SPR experiments binding of Staufen dsRBD3-4-L4 to surface-coupled Tm1 FL was observed.** The overlaid sensorgrams of a multi-cycle SPR experiment (left) show that binding occurs transiently. $K_D$s were determined using a one-site-binding fit to the maximal response at equilibrium for each analyte concentration. The $K_D$-value is given as mean ± SD from triplicate experiments.

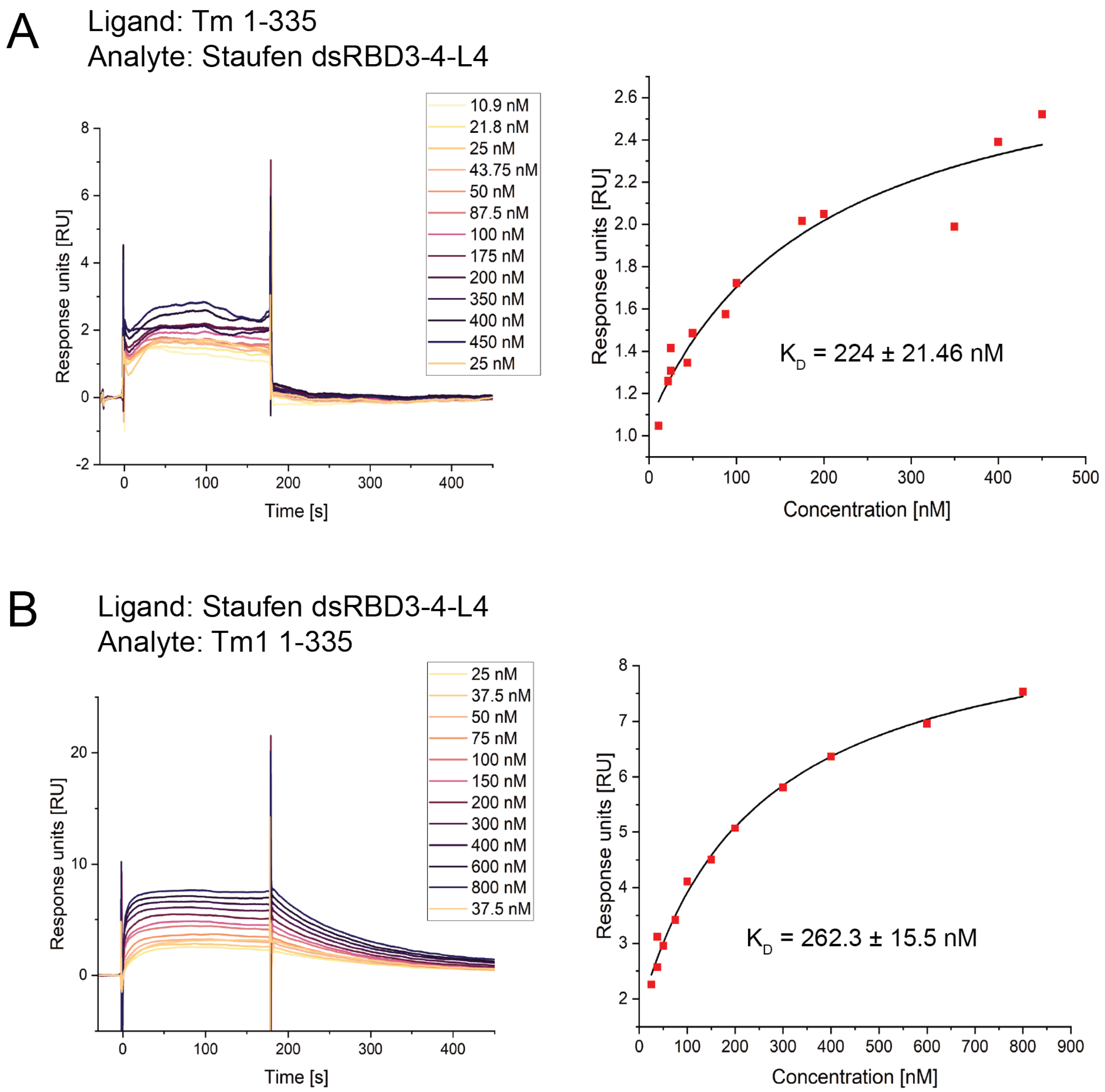


**Figure S5: In SPR experiments an interaction between Staufen dsRBD3-4-L4 and Tm1 1-335 is observed. A-B)** Multi-cycle SPR experiments show binding of Staufen dsRBD3-4-L4 to surface-coupled Tm1 1-335 (A) and Tm1 1-335 to surface-coupled Staufen dsRBD3-4-L4 (B). The overlaid sensorgrams of multi-cycle SPR experiments (left) show that binding occurs transiently. $K_D$ was determined using a one-site-binding fit to the maximal response at equilibrium for each analyte concentration (right). The $K_D$-values are given as mean ± SD from triplicate experiments.

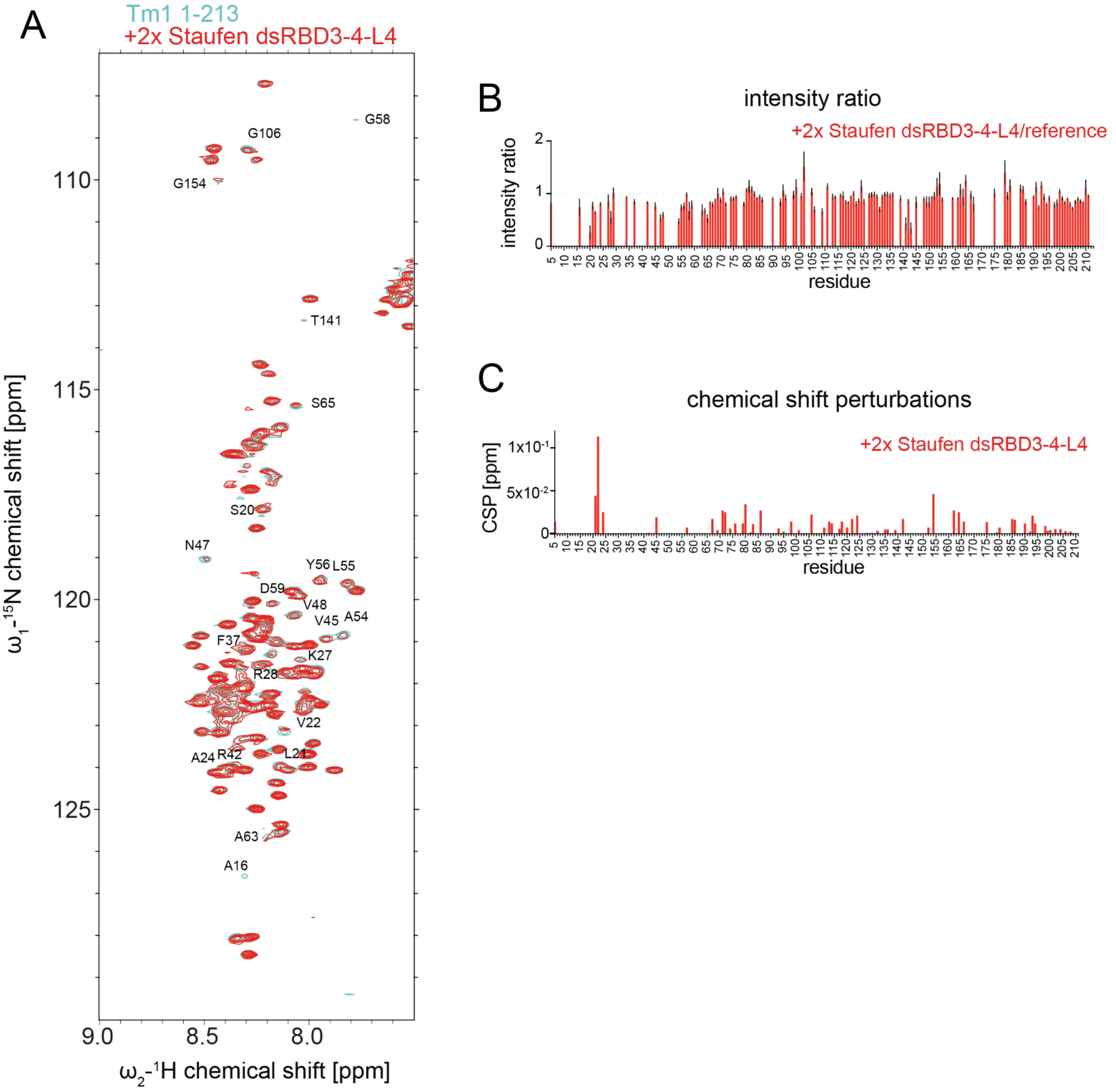


**Figure S6: $^{1}$H-$^{15}$N HSQC titration experiment with Tm1 1-213 and Staufen dsRBD3-4-L4. A)** $^{1}$H-$^{15}$N HSQC spectrum of Tm1 1-213 with backbone assignments indicated (blue), overlaid with its $^{1}$H-$^{15}$N HSQC spectrum after titration with a 2-fold excess of Staufen dsRBD3-4. **B)** Plotted intensity ratios and **C)** chemical shift perturbation (CSP) plot of individual residues in the titration experiment. Data are presented as measured value ± SD. Error bars represent estimates of propagated measurement errors of the experimental uncertainties in signal amplitudes.

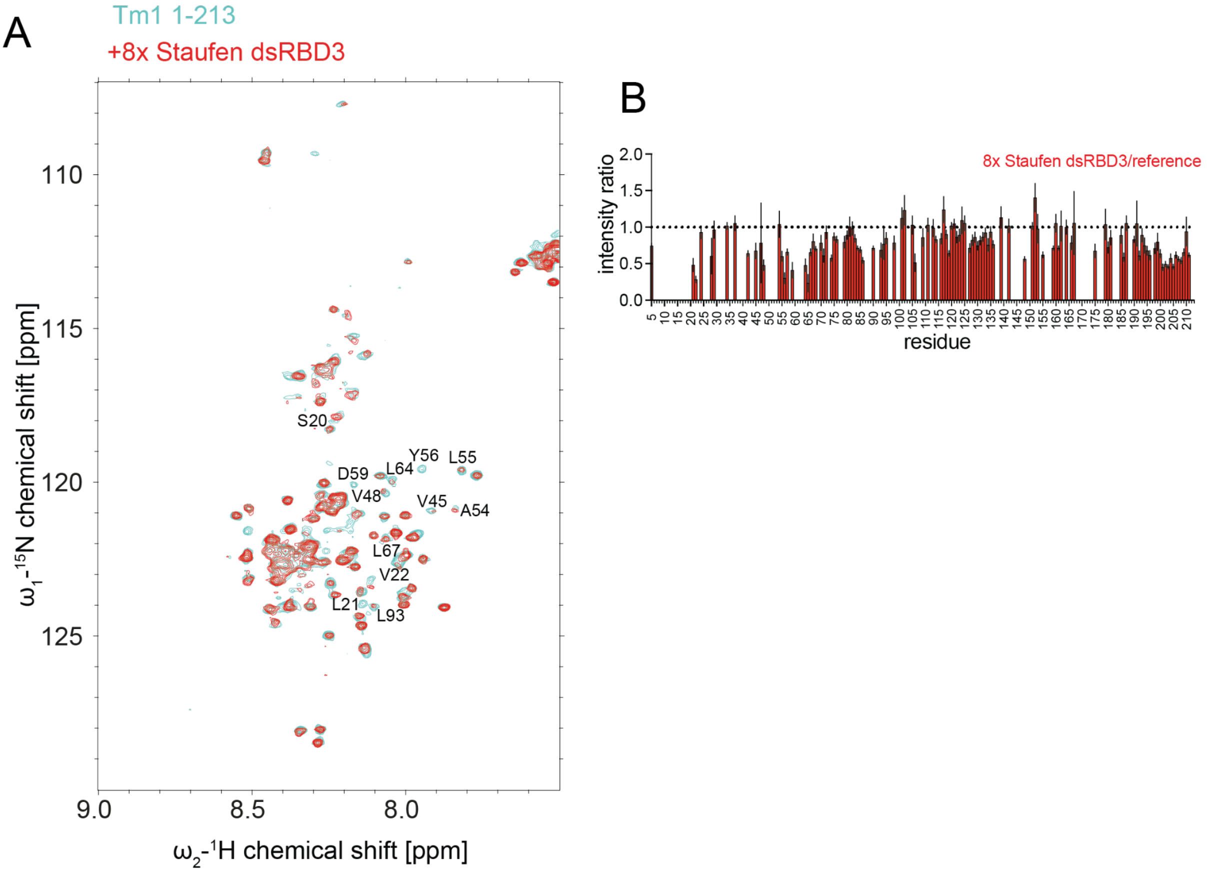


**Figure S7: $^{1}$H-$^{15}$N HSQC titration experiment with Tm1 1-213 and Staufen dsRBD3. A)** $^{1}$H-$^{15}$N HSQC spectrum of Tm1 1-213 with backbone assignments (Vaishali *et al*, 2021) indicated (cyan), overlaid with its $^{1}$H-$^{15}$N HSQC spectrum after titration with an 8-fold excess of Staufen dsRBD3 (red). **B)** Plotted intensity ratios of individual residues in the titration experiment. Data are presented as measured value ± SD. Error bars represent estimates of propagated measurement errors of the experimental uncertainties in signal amplitudes.

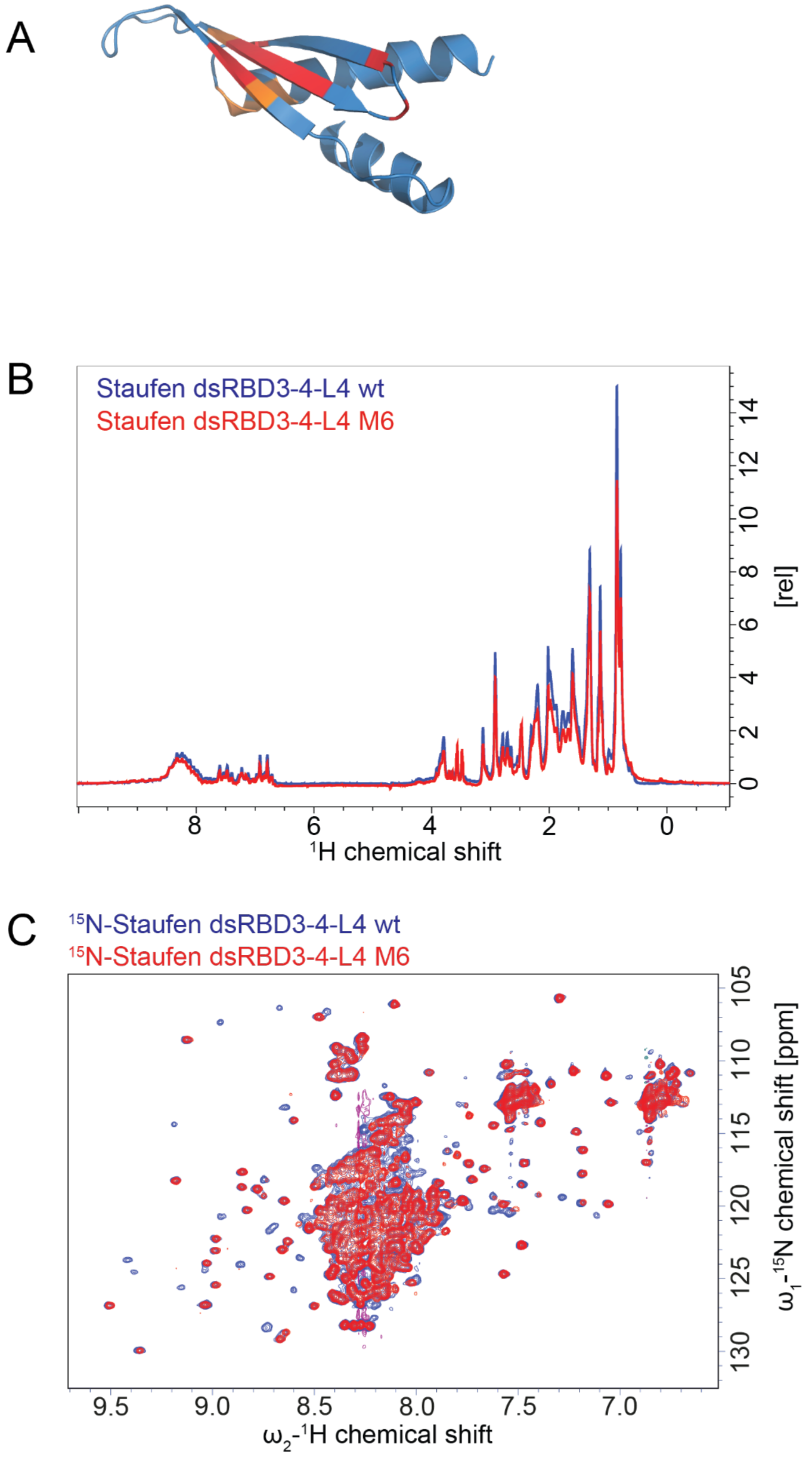


**Figure S8: The Staufen M6 mutant protein retains wildtype-like folding. A)** AlphaFold2 (Jumper *et al*, 2021) prediction of the Staufen dsRBD4 structure with RNA-binding residues marked in orange and Tm1-binding residues marked in red. **B-C)** Overlay of 1D (A) and $^{1}H$-$^{15}N$ HSQC (B) spectra of Staufen dsRBD3-4-L4 WT (blue) and the mutant M6 (red).

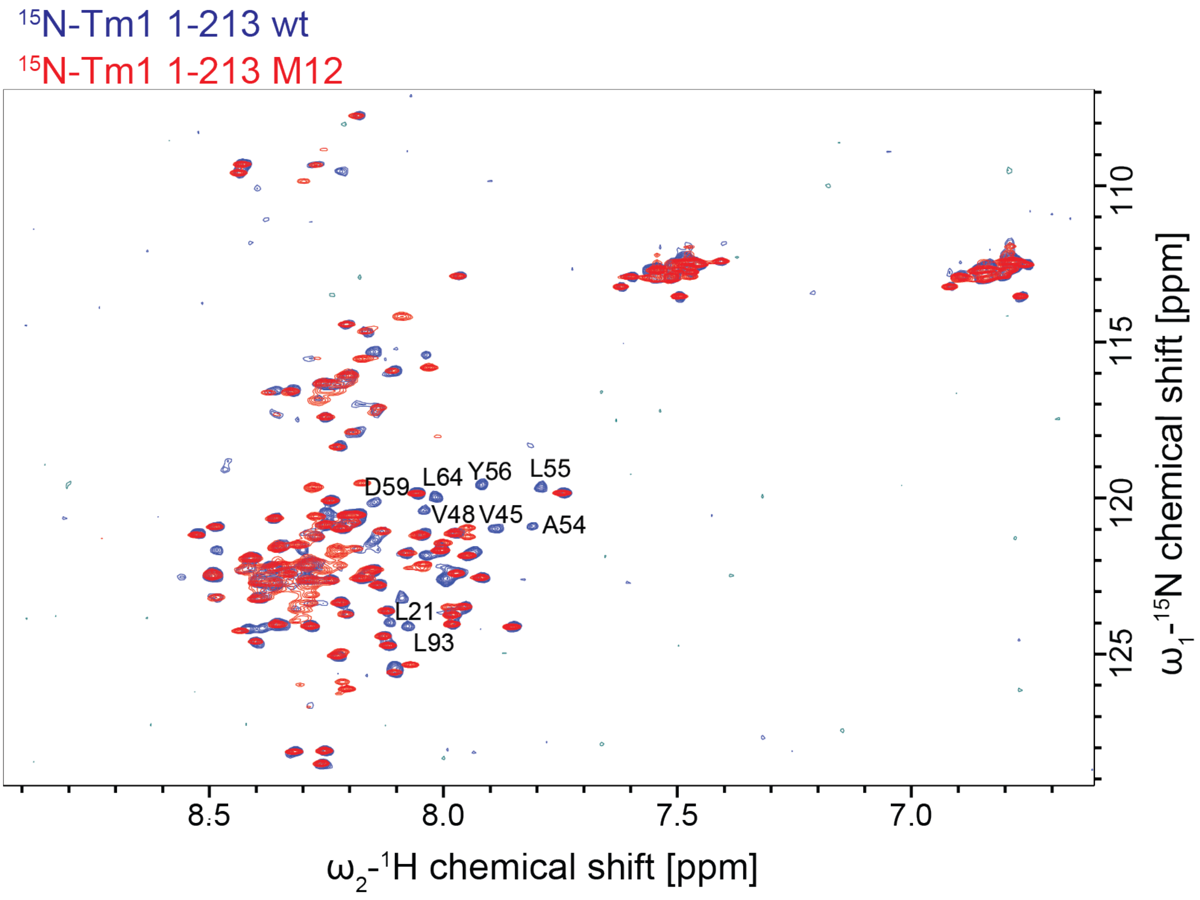


**Figure S9: The Tm1 M12 mutant protein remains unfolded.** Overlay of $^{1}H$-$^{15}N$ HSQC spectra of Tm1 1-213 WT (blue) and the mutant M12 (red).

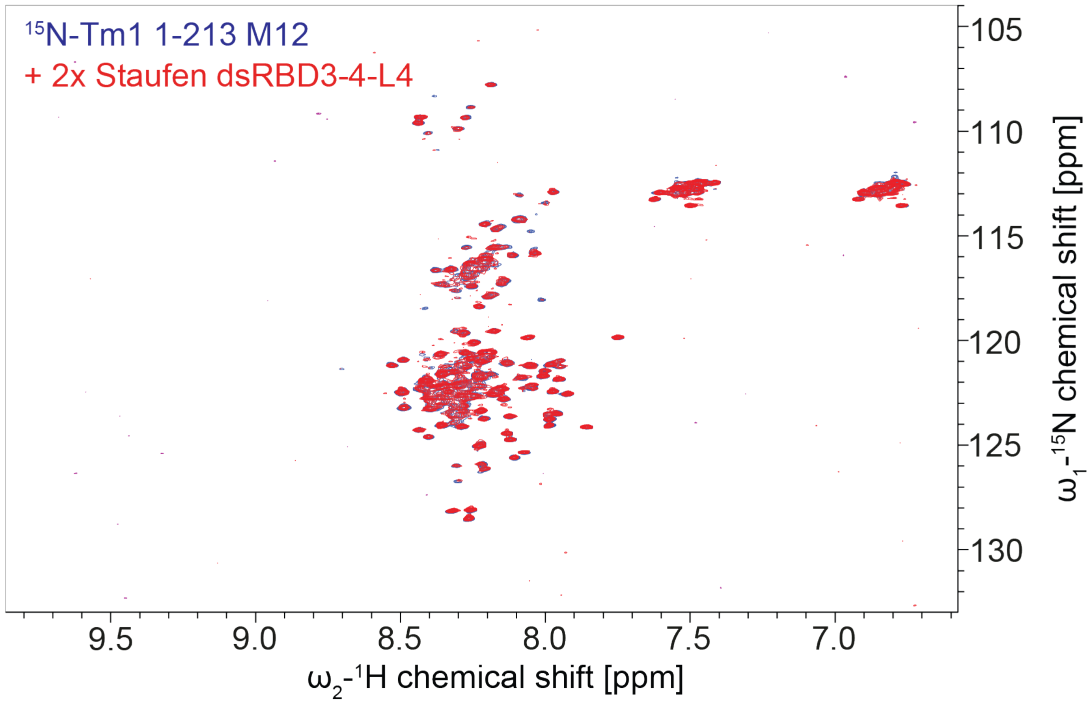


**Figure S10: Tm1 1-213 M12 does not interact with Staufen dsRBD3-4-L4 at micromolar concentrations.** $^{1}$H-$^{15}$N HSQC titration experiment with 50 µM $^{15}$N-labeled Tm1 1-213 and Staufen dsRBD3-4-L4 showed no changes of the Tm1 1-213 upon addition of Staufen dsRBD3-4-L4.

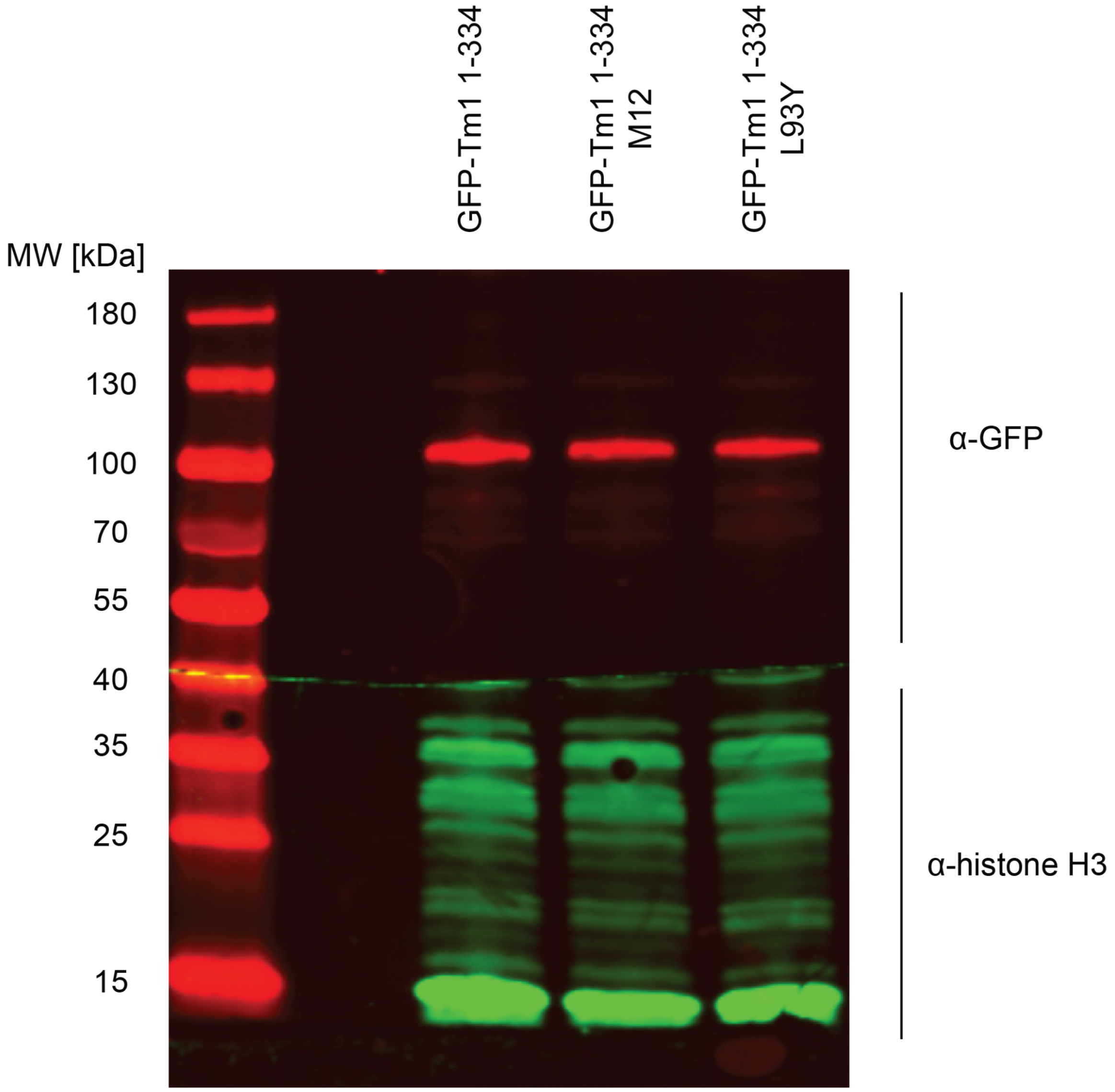


58 **Figure S11:** Western blot analysis of GFP-Tm1 proteins expressed from transgenes in
59 *Tm1[eg9]/Tm1[eg9] Drosophila* ovaries. All transgenes are expressed to similar levels from a UAS
60 promoter, driven by *oskar-Gal4*.

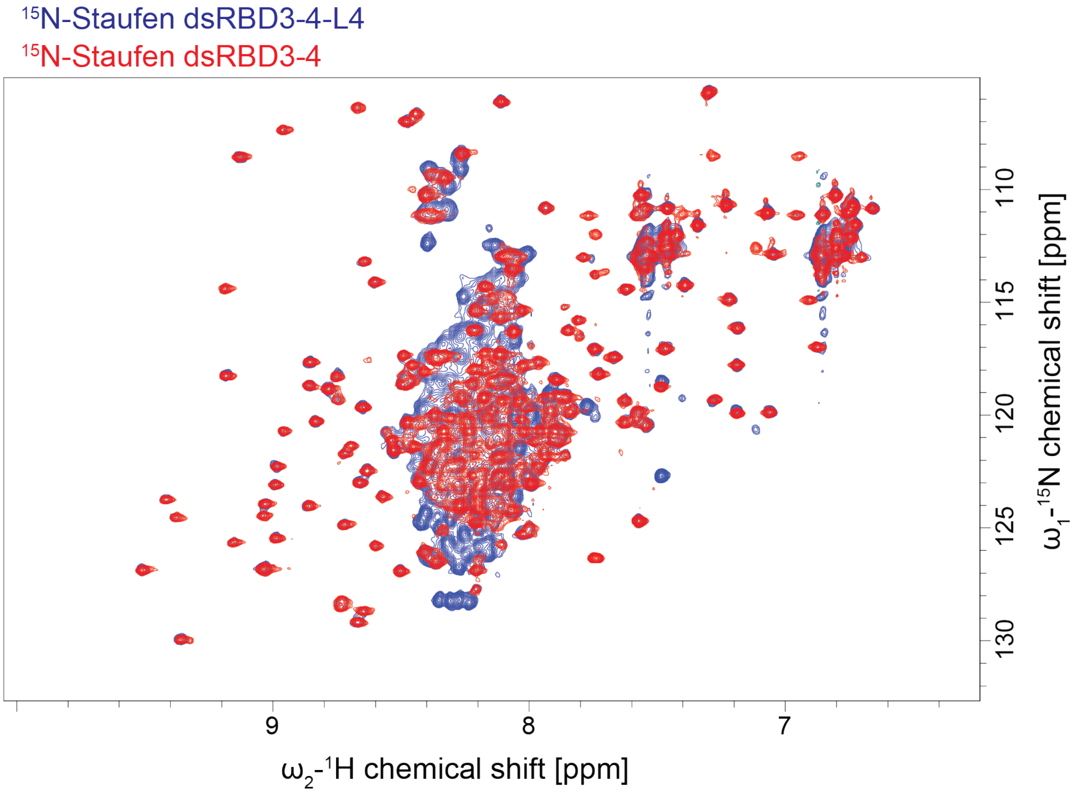


**Figure S12: Truncation of linker 4 does not alter the fold of Staufen dsRBDs 3 and 4.** [1] $^{1}$H-$^{15}$N HSQC spectrum of Staufen dsRBD3-4-L4 (blue), overlaid with the $^{1}$H-$^{15}$N HSQC spectrum of Staufen dsRBD3-4 (red).

**Table S1:** Overview of all Staufen and Tm1 proteins, constructs, and mutants used in this work. $K_D$ of their interaction with their respective interaction partner as determined by SPR are indicated.

| Staufen protein construct | Binding to Tm1 | $K_D$ |
|---|---|---|
| Staufen FL (aa 1-1026) | yes | 86.6 nM ± 17.4 nM |
| Staufen Nterm-dsRBD1-2 (aa 1-557) | no | - |
| Staufen dsRBD3-4-5-Cterm (aa 578-1026) | yes | 152 nM ± 60.01 nM |
| Staufen dsRBD1-4 (aa 311-781) | no | - |
| Staufen L4-dsRBD5-Cterm (aa 782-1026) | no | - |
| Staufen dsRBD5-Cterm (aa 951-1026) | no | - |
| Staufen dsRBD3-4-L4 (aa578-950) | yes | 224 nM ± 21.46 nM |
| Staufen dsRBD3-4-L4 M6 (aa578-950 RTY FSV) | no | - |
| **Tm1 protein construct** | **Binding to Staufen** | **$K_D$** |
| Tm1 FL (aa 1-441) | yes | 86.6 nM ± 17.4 nM |
| Tm1 1-335 | yes | 385.9 nM ± 44.8 nM |
| Tm1 1-213 | yes | 494.8 nM ± 89.3 nM |

| | | |
|---|---|---|
| Tm1 1-213 M12 | no | - |
| Tm1 1-213 M3 | yes | 976.4 nM -± 85.6 nM |
| Tm1 1-213 M8 | yes | 1246.3 nM -± 94.9 nM |
| Tm1 1-213 L93Y | yes | 1421 nM -± 143.5 nM |
